# IL-18 produced by pregnant uterus promotes essential inflammatory responses and fetoplacental growth

**DOI:** 10.1101/2025.04.23.649615

**Authors:** Hajime Ino, Yasuyuki Negishi, Yumi Horii, Eri Koike, Richard A. Flavell, Shunji Suzuki, Rimpei Morita

## Abstract

Placental insufficiency affects fetomaternal health throughout life. Although the interaction between the maternal uterine immune milieu and fetal-derived cells plays a crucial role in placental formation, several aspects remain unclear. Therefore, we conducted this study to investigate the effects of interleukin (IL)- 18, a unique cytokine with both proinflammatory and anti-inflammatory properties, on the uterine immune milieu and placental development. Our results identified pregnant uterine smooth muscle cells as an important source of IL-18, which supports homeostatic type 1 immune responses. IL-18 facilitates appropriate placental development through uterine vascular remodeling and placental angiogenesis. Smooth muscle cell–specific *Il18*-knockout dam mice exhibited impaired fetoplacental growth and elevated maternal blood pressure, reflecting preeclampsia-like phenotypes. Their offspring demonstrated a tendency toward excessive weight gain and delayed neurodevelopment. Overall, this study emphasizes the essential role of IL-18 in placental formation and its wider implications for fetomaternal health.

**Graphical Abstract:** 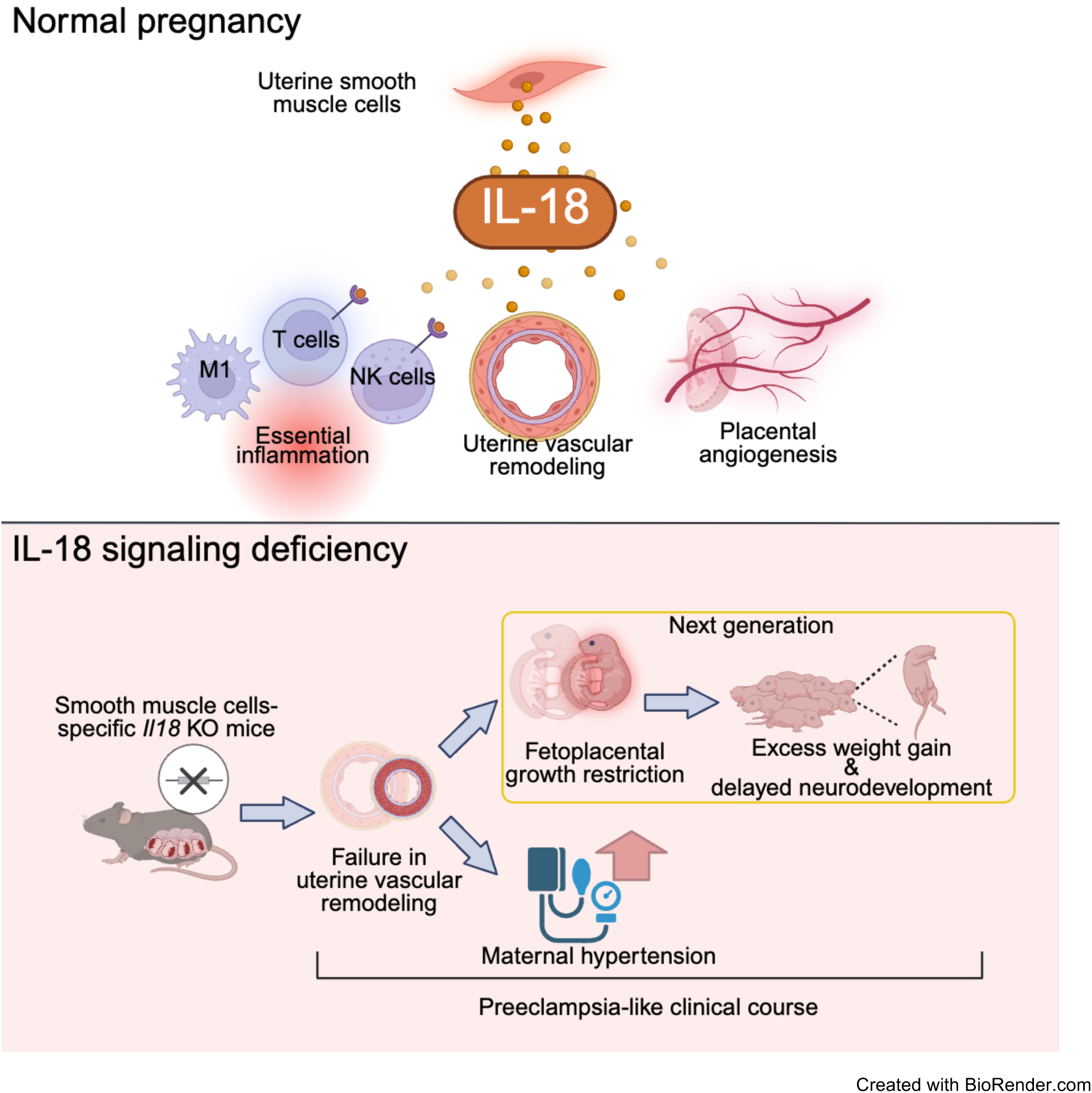

**Highlights:**

- Smooth muscle cells in the pregnant uterine myometrium sufficiently produce IL-18
- IL-18 induces type 1 immune responses in the uterine immune milieu
- IL-18 promotes maternal uterine vascular remodeling and placental angiogenesis
- IL-18-signaling deficiency in the mouse model shows the preeclampsia-like clinical course

## Introduction

Formation of the placenta in mammals is a fundamental process for species survival. However, placental insufficiency has been implicated in maternal multiple organ failure and preeclampsia ^1,2^ and can result in long-term health complications in subsequent generations ^3–5^. This implies that revealing the hidden mechanisms of placental formation can contribute to the lifelong health of both the mother and child. During pregnancy, the immune environment undergoes remarkable changes across each gestational stage. Although excessive inflammation exerts detrimental effects in all trimesters, an appropriate inflammatory response is essential during periods of maternal tissue remodeling that is required for successful implantation and placental growth ^6,7^. Hence, this dynamic immunological environment emphasizes the importance of understanding the roles of inflammatory cytokines in placental formation.

Interleukin (IL)-18 is a member of the IL-1 family cytokines, which is produced as a 24-kDa precursor and is cleaved into an 18-kDa mature form by caspase-1 that is activated by the inflammasome ^8^. IL-18 forms a complex with IL-18 receptor α (Rα) and β (Rβ) chains at the plasma membrane, which signals into the cell ^9^. IL-18 is potentially produced by various immune and non-immune cells, although it is generally produced by monocytic cells such as macrophages, which transmit its signal into T cells and NK cells where IL-18 receptors (IL-18Rs) are highly distributed ^10^. Among the family cytokines, IL-1β possesses a strong proinflammatory property and exerts adverse effects on pregnancy, including preterm birth ^11,12^, preeclampsia ^13^, and fetal neurodevelopmental disorders ^14^. In contrast, IL-18, which shares a similar production pathway with IL-1β, possesses both proinflammatory and anti-inflammatory properties. IL-18 is implicated in tissue and vascular remodeling in inflammatory airway diseases ^15,16^ and rheumatoid arthritis ^17^, partially due to its unique duality; however, the role of IL-18 in pregnancy remains largely unclear.

The uterine decidua is the immune surface where fetus-derived extravillous trophoblasts and mother- derived immune cells cross-talk and has long been considered the reproductive immune microenvironment that exerts the greatest impact on pregnancy outcomes ^18,19^. Nevertheless, it has been reported that a specific subtype of smooth muscle cells (SMCs) expressing interferon (IFN)-γ exists in the human uterine myometrium and is involved in parturition ^20^, and IL-33 produced by myofibroblasts and other cells in the myometrium promotes remodeling of the maternal uterine spiral artery through type 2 immune responses^21^. The integration of recent and previous findings suggests that all immune and nonimmune cells throughout the uterine organ potentially produce cytokines, which regulate homeostatic pregnancy processes, including the establishment, maintenance, and termination of pregnancy.

We conducted this study to investigate the impact of homeostatically produced IL-18 on the uterine immune milieu and fetoplacental development using mouse models. We first identified the major source of IL-18 in pregnant mice and then explored its effects on the uterine immune milieu through both *in vivo* and *in vitro* experiments by administering an IL-18-neutralizing antibody (IL-18 nAb). Our results showed that uterine myometrial SMCs are a vital source of IL-18. Next, using SMC-specific *Il18*-knockout mice, we investigated the effects on upstream and downstream signaling pathways using RNA sequencing (RNA-seq), fetoplacental development, and the growth and neurodevelopment of subsequent generations. This study clarifies the correlation between the uterine immune environment and fetoplacental development centered on IL-18, and moreover, through the novel reproductive immunological approach for understanding placental development, this study will provide strategies for the early detection and prevention of adverse pregnancy outcomes such as preeclampsia.

## Results

### SMCs in the pregnant uterine myometrium sufficiently produce IL-18

Tissue lysates derived from the uterine myometrium, uterine decidua, and placenta were analyzed by western blotting to identify the organs responsible for IL-18 production in pregnant mice. In the pregnant myometrium, IL-18 precursor and its cleaved form were sufficiently detected without additional stimulation (Figure 1A). Moreover, compared with the decidua and placenta, the myometrium contained abundant levels of molecules, such as caspase-1, NLRP3 and gasdermin-D, which are associated with inflammasome activation—a canonical pathway that mediates IL-18 production (Figures 1A, S1A). Next, multicolor flow cytometry was used to identify the cells that produce IL-18 in the pregnant uterine myometrium. Although a wide variety of immune and non-immune cells can produce IL-18, macrophages are among the most prominent sources of this cytokine ^10^. In fact, macrophages in the uterine myometrium were an important source of IL-18, although CD45-negative non-immune cells also produce substantial amounts of IL-18 (Figures 1B, S1B). Considering the large proportion of these CD45-negative cells in the tissue, we concluded that they are the most significant source of IL-18 in the pregnant myometrium (Figure 1B). The majority of these CD45-negative cells were αSMA-positive and identified as SMCs (Figure S1B). These SMCs exhibited abundant intracellular IL-18 production compared with other immune cells (Figure

**Figure 1.**
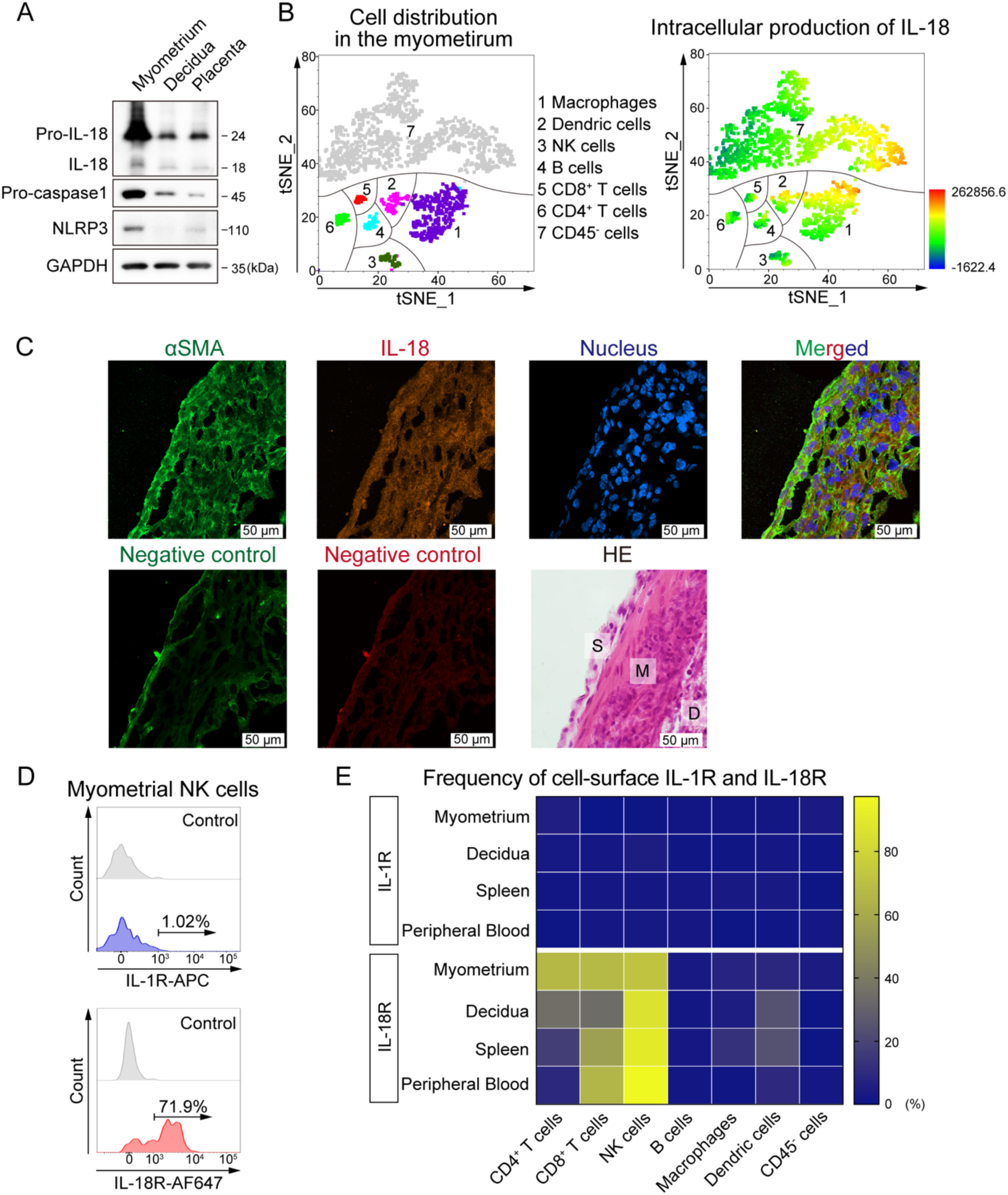
SMCs in the pregnant uterine myometrium abundantly produce IL-18. (A) Immunoblotting using the indicated antibodies. The uterine myometrium, decidua, and placenta were extracted from pregnant mice, and cells were collected from whole tissue lysates. The image is representative of two independent samples. (B) *t*-distribution stochastic neighbor embedding (t-SNE) visualization of uterine myometrial cells extracted from pregnant mice at E13.5. Cell distribution (left) and intracellular production of IL-18 (right) were determined by flow cytometry. IL-18 signal intensities are depicted according to the rainbow lookup bar. The image is representative of two independent samples. (C) Confocal fluorescence immunohistochemistry images using the indicated antibodies. Pregnant uteri were extracted at E9.5. Negative control (stained with secondary antibody only, without a primary antibody) and HE staining of proximal sections are also shown. The HE-stained image also shows the position of each tissue (S: uterine serosa, M: uterine myometrium, D: uterine decidua). Each image is representative of three independent samples. (D) Representative histogram of flow cytometric analysis depicting allophycocyanin (APC)-labeled cell surface IL-1R and IL-18R in uterine myometrial NK cells collected from pregnant mice. Each image is representative of two independent samples. (E) Heatmap generated from flow cytometric analysis of the uterine myometrium, decidua, spleen, and peripheral blood collected from pregnant mice shows the frequencies of the cell-surface IL-1R and IL- 18R on each cell. The frequency defining the heatmap was calculated using the average of two independent samples. Related to Figure S1.

S1B), and fluorescence immunostaining of frozen sections of placenta-implantation sites also demonstrated abundant IL-18 production in SMCs, which are abundant in the myometrium (Figure 1C). Although there are reports of IL-18 production by bronchial ^22^ and vascular ^23^ SMCs, this is the first report of IL-18 production by uterine SMCs. Furthermore, myometrial SMCs expressed NLRP3 and caspase-1, which are factors essential for canonical inflammasome activation in the IL-18 production process (Figures S1C, S1D). Activation of noncanonical caspases, including caspase-8 ^24^ and caspase-11 ^25^, was also detected in the myometrium (Figure S1A), indicating that IL-18 production in the pregnant uterus is not limited to a single pathway.

Next, the distribution of IL-18Rs in pregnant mice was evaluated by flow cytometry. IL-18Rs were particularly highly distributed on T cells and NK cells in the pregnant uterus (Figures 1D, 1E). Moreover, IL-18Rs showed higher expression levels than IL-1 receptor (IL-1R; the receptor for IL-1β, another member of the IL-1 superfamily) (Figures 1D, 1E, S1E); this high expression was stably maintained throughout pregnancy (Figure S1E).

Altogether, our results identified uterine SMCs in the pregnant myometrium as a critical source of IL-18. In addition, both canonical and noncanonical pathways may be involved in IL-18 production. Finally, our results showed that IL-18Rs is abundant on T cells and NK cells in the pregnant uterus, suggesting a crucial role for IL-18-signaling.

### IL-18 promotes type 1 immune responses in the uterus *in vivo*

We investigated the effect of IL-18 on the uterine immune environment *in vivo* for which pregnant mice were repeatedly administered an IL-18 nAb or its isotype IgG, and cells collected from the excised uterine myometrium and decidua were analyzed by flow cytometry (Figures 2A, S2A). According to previous reports, IL-18 promotes the production of the inflammatory cytokine IFN-γ by T cells and other cells via a type 1 immune response in the presence of IL-12, whereas in the absence of IL-12, it promotes the production of the anti-inflammatory cytokine IL-4 via a type 2 immune response ^26^. Moreover, IFN-γ is essential for the remodeling of uterine spiral arteries and angiogenesis at the implantation site ^27^, and IL- 4 contributes to the maintenance of pregnancy by suppressing excessive inflammation ^28^. Therefore, we investigated the intracellular production levels of IFN-γ, IL-4, and IL-12 to determine how IL-18 orchestrates the balance of the uterine immune environment.

**Figure 2.**
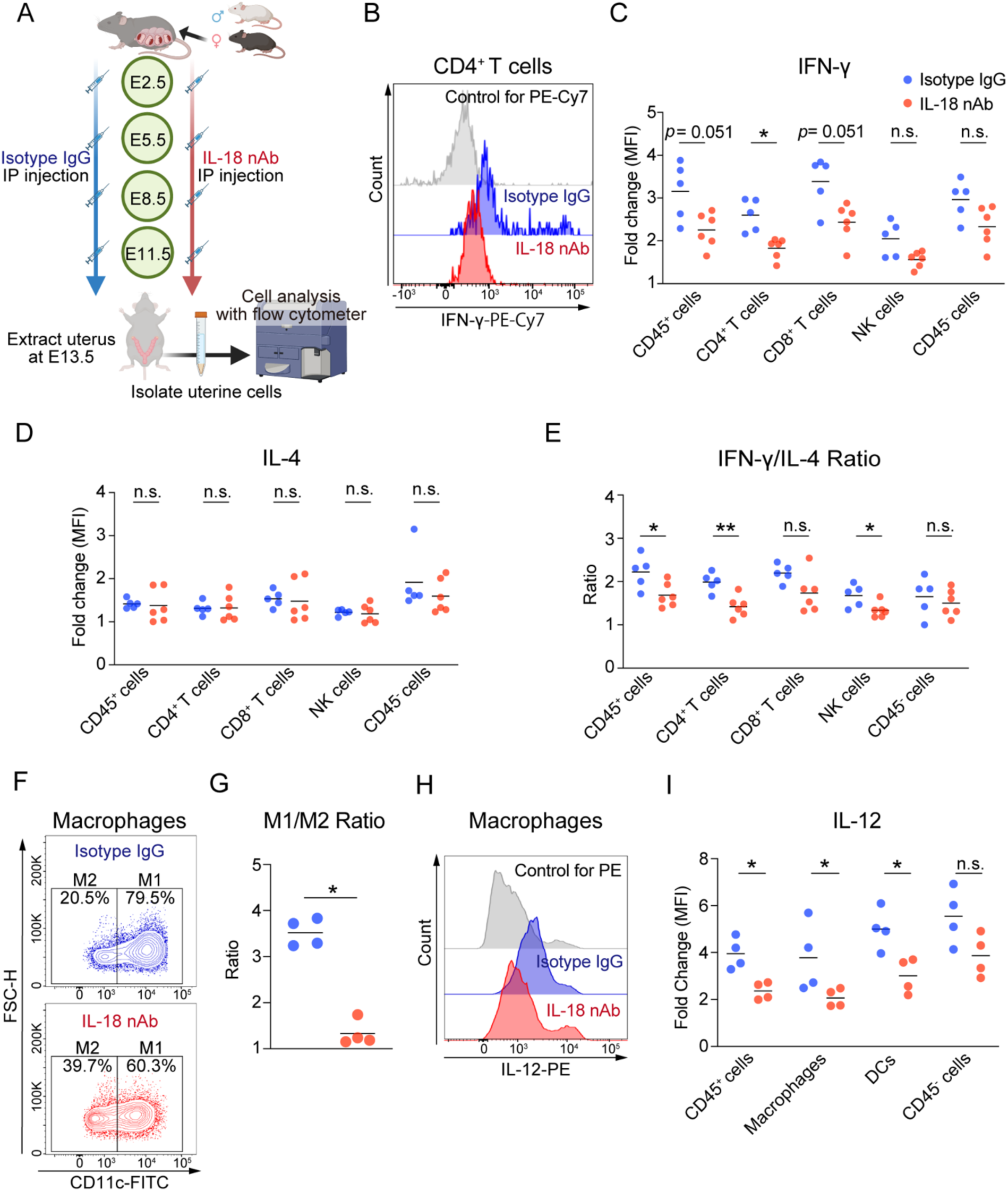
IL-18 promotes type 1 immune responses in the uterus *in vivo*. (A) B6 WT female mice were mated with Balb/c WT male mice and injected intraperitoneally with 600 µg of anti-IL-18-neutralizing antibody (IL-18 nAb) or 600 µg of its isotype IgG at E2.5, E5.5, E8.5, and E11.5. At E13.5, the pregnant uterine myometria were extracted, and cells were isolated and analyzed by flow cytometry. Created with BioRender.com. (B) Representative histogram of flow cytometric analysis depicting phycoerythrin-cyanine7 (PE-Cy7)- labeled intercellular interferon gamma (IFN-γ) in myometrial CD4^+^ T cells by group. (C-D) Mean fluorescence intensity (MFI) of PE-Cy7-labeled intracellular cytokines in each myometrial immune cell population is shown for relative change to control by group (Mann–Whitney U test; each data point represents 1 dam, n = 5–6 per each group, each horizontal line represents the mean value of samples). (E) Ratio of intracellular IFN-γ/IL-4 MFI in each immune cell is shown by group (Mann–Whitney U test, each data point represents 1 dam, n = 5–6 per each group, each horizontal line represents the mean value of samples). (F) Representative contour plots of flow cytometric analysis showing the distribution of M1 (F4/80^+^CD11c^+^) and M2 (F4/80^+^CD11c^-^) macrophages identified using fluorescein isothiocyanate (FITC)-labeled CD11c. (G) Ratios of M1/M2 cell population in macrophages are shown by group (Student’s *t*-test, each data point represents 1 dam, n = 4 per each group, each horizontal line represents the mean value of samples). (H) Representative histogram of flow cytometric analysis depicting phycoerythrin (PE)-labeled intercellular IL-12 in uterine macrophages by group. (I) MFI of PE-labeled intracellular IL-12 in each immune cell population is shown for relative change to control by group (Mann–Whitney U test, each data point represents 1 dam, n = 4 per each group, each horizontal line represents the mean value of samples). * *p* < 0.05, ** *p* < 0.01 Related to Figure S2.

Flow cytometric analysis revealed no differences in the proportions and numbers of CD4^+^ T cells, CD8^+^ T cells, and NK cells in the uterine myometrium between the isotype IgG group and IL-18 nAb group (Figure S2B). Compared with the isotype IgG group, the IL-18 nAb group demonstrated a tendency of reduced intracellular IFN-γ production in myometrial CD4^+^ T cells, CD8^+^ T cells, and NK cells, whereas IL- 4 production showed no significant alterations (Figures 2B, 2C, 2D, 2E). T cells and NK cells in the decidua exhibited a similar trend (Figure S2C). These data suggest that IL-18–IL-18R signaling promotes a type 1 immune response mediated by T cells and NK cells throughout the uterus.

We next examined the effect of IL-18 on antigen-presenting cells. After classifying uterine myometrial macrophages into M1 (F4/80^+^CD11c^+^) and M2 (F4/80^+^CD11c^−^) subtypes, we detected an increase in the proportion of M2 macrophages in the IL-18 nAb group compared with that in the isotype IgG group (Figures 2F, 2G), suggesting that IL-18 helps maintain a predominant M1 state in macrophages. In contrast, when dendritic cells (DCs) were subdivided into DC1 (F4/80^−^CD11c^+^CD103^+^) and DC2 (F4/80^−^CD11c^+^CD103^−^), we found no significant difference between the two groups (Figures S2D, S2E). Because antigen-presenting cells are the major source of IL-12, we then investigated intracellular IL-12 production in both macrophages and DCs and observed a significant reduction in the IL-18 nAb group compared with that in the isotype IgG group (Figures 2H, 2I). Although antigen-presenting cells exhibit a lower positivity of IL- 18R expression than other immune cells (Figure 1E), suggesting that IL-18 exerts these effects indirectly, our results indicate that IL-18 promotes M1 polarization in macrophages and IL-12 production in antigen-presenting cells.

Overall, our *in vivo* experiments demonstrated that IL-18–IL-18R signaling comprehensively improves type 1 immune responses in the uterus by inducing IFN-γ production by T cells and NK cells, as well as IL-12 production by antigen-presenting cells.

### IL-18 promotes both IFN-γ and IL-4 production in the uterine myometrium *in vitro*

To confirm the phenomena observed *in vivo*, we designed an *in vitro* model of the pregnant uterine immune milieu. Hematopoietic (CD45^+^) cells were enriched from embryonic day 13.5 (E13.5) pregnant uteri using magnetic-activated cell sorting (MACS) and then incubated overnight with either IL-18 nAb or its isotype IgG. This was followed by flow cytometric analysis of the cells and protein analysis of the culture supernatant by enzyme-linked immunosorbent assay (ELISA) (Figure 3A). No pharmacological agents were added to stimulate the cells in this experimental setup. The enrichment of CD45^+^ cells by MACS achieved a purity of >80% (Figure S3A).

**Figure 3.**
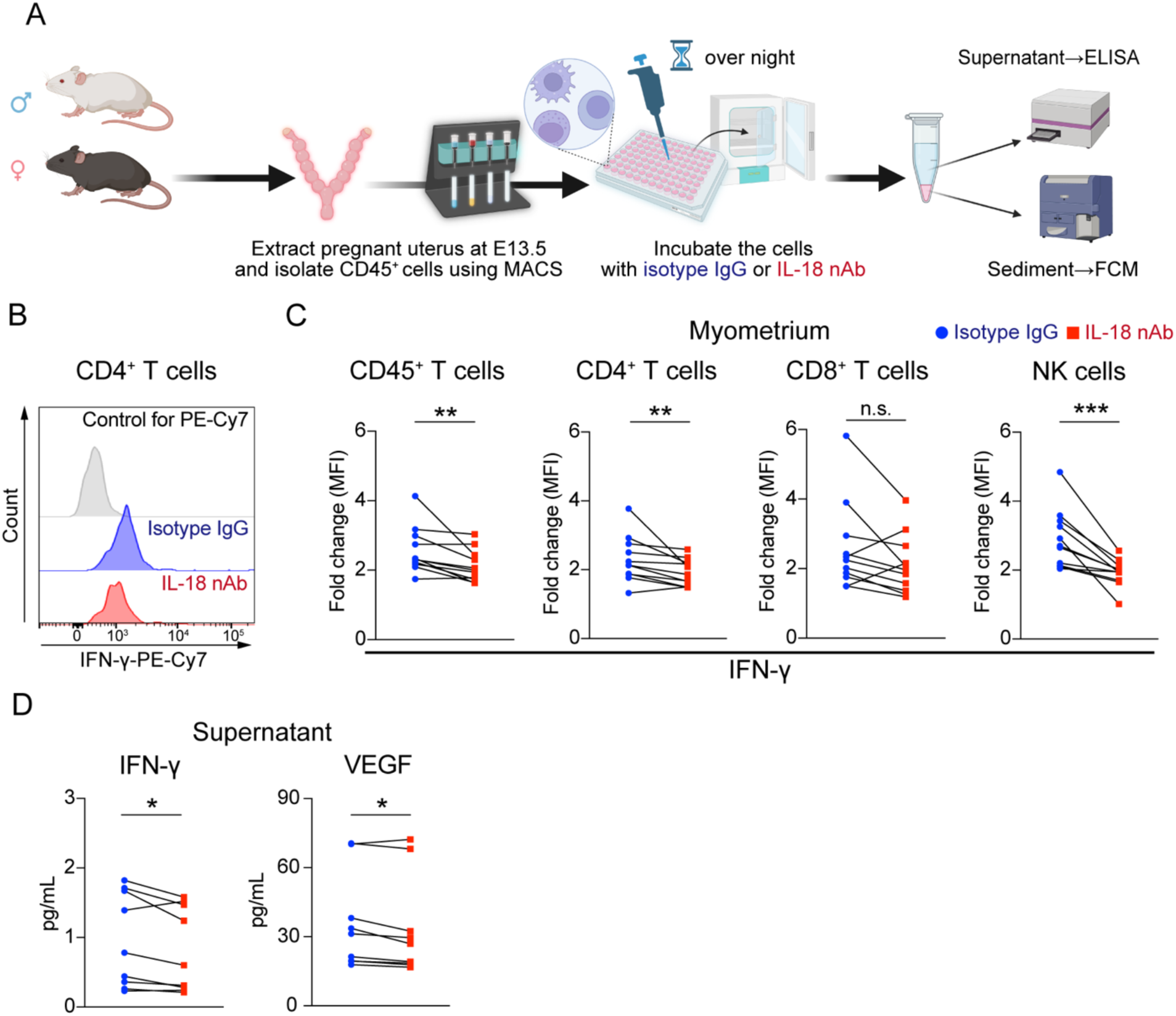
IL-18 promotes IFN-γ production in uterine immune milieu *in vitro*. (A) B6 WT female mice were mated with Balb/c WT male mice, and pregnant uteri were extracted at E13.5. CD45^+^ cells were isolated by magnetic-activated cell sorting (MACS) from the myometrium or decidua, and the cells were incubated with IL-18 nAb or its isotype IgG. The cells were then analyzed by flow cytometry, and the supernatants were subjected to ELISA. Created with BioRender.com. (B) Representative histogram of flow cytometric analysis depicting PE-Cy7-labeled IFN-γ in myometrial CD4^+^ T cells by group. (C) MFI of PE-Cy7-labeled intracellular IFN-γ in each myometrial immune cell population is shown for relative change to control by group (Wilcoxon matched-pairs signed rank test, each pair of data points represents 1 dam, n = 11 per each group). (D) Concentration of IFN-γ and VEGF in the culture supernatant quantified by ELISA is shown (Wilcoxon matched-pairs signed rank test, each pair of data points represents 1 dam, n = 9 per each group). * *p* < 0.05, ** *p* < 0.01, *** *p* < 0.001 Related to Figure S3.

Consistent with our *in vivo* findings, the administration of IL-18 nAb reduced intracellular IFN-γ production in myometrial T cells and NK cells compared with that observed by the administration of the isotype IgG (Figures 3B, 3C). Moreover, the IFN-γ concentration in the culture supernatant was lower in the IL-18 nAb group than in the isotype IgG group. Considering that a previous study reported that IL-18 promotes VEGF production by synovial fibroblasts in rheumatoid arthritis ^29^, we measured VEGF levels in the culture supernatant and found them to be significantly reduced in the IL-18 nAb group compared with those in the isotype IgG group (Figure 3D). In contrast to the *in vivo* results, the administration of IL-18 nAb also reduced intracellular IL-4 production in myometrial CD4^+^ T cells compared with that in the isotype IgG group (Figures S3B, S3C). However, considering that IL-4 concentration in most culture supernatant samples was less than the detection sensitivity (Figure S3D), the absolute amount of IL-4 produced by immune cells in the pregnant uterus may be extremely low. The IL-12 levels in the culture supernatant showed no significant difference between the two groups (Figure S3D). Despite the limited sample size, similar experiments conducted using cells derived from the uterine decidua revealed no significant differences in intracellular IFN-γ or IL-4 production between the two groups (Figure S3E).

Collectively, these findings confirm that IL-18 promotes IFN-γ production by T cells and NK cells in the myometrium *in vitro*, consistent with the *in vivo* findings.

### IL-18 indicates to support placental development and fetal growth

Based on previous reports that IL-18 promotes tissue remodeling in chronic inflammatory diseases ^15,30^ and that IL-33, another IL-1 superfamily cytokine, facilitates fetal growth through uterine vascular remodeling ^21^, we hypothesized that IL-18 supports fetal growth through remodeling maternal tissues. To test this hypothesis, we conducted an experiment in which pregnant mice were repeatedly injected with either IL-18 nAb or its isotype IgG control, after which the body weight of newborn pups was measured. Pups born to IL-18 nAb-treated dams had significantly lower birth weights than those born to mice in the control group (Figure S4A). Furthermore, to confirm the importance of IL-18-mediated type 1 immune responses, we compared neonatal body weights between wild-type (WT) and *Ifng* (encoding IFN-γ protein) total knockout (KO) pregnant mice. As anticipated, pups born to *Ifng* KO dams showed significantly lower birth weights (Figure S4B). These data suggest that IL-18-signaling and type 1 immune responses play vital roles in fetal growth.

We next performed a structural analysis of the placenta to determine whether fetal growth restriction (FGR) due to IL-18-signaling deficiency was attributable to impaired placental development. Our results showed that placentas derived from IL-18 nAb-treated pregnant mice exhibited a poorly developed vascular structure and an atrophic parenchyma in the placental labyrinth resembling the villous structures of the human placenta (Figures S4C, S4D). This observation was further confirmed by immunofluorescence staining of CD31^+^ endothelial cells, which revealed a similarly defective vascular complexity (Figure S4E). These results suggest that the placental labyrinth remained immature, causing insufficient vascular complexity and a reduced efficiency of nutrient and oxygen exchange at the maternal– fetal interface ^31,32^. Hence, our findings indicate that IL-18 plays a vital role in placental function and fetal development.

### SMC-specific *Il18* KO mice exhibit a preeclampsia-FGR-like clinical course

By focusing on the finding that uterine SMCs are a vital source of IL-18, we next established SMC-specific IL-18 conditional knockout mice (*Il18^fl/fl^;Sm22a-cre*) using the Cre-loxP system with *Il18^fl/fl^* mice as controls. After mating these dams with wild-type male mice, we conducted the following assessments: morphological evaluation of spiral artery remodeling in excised uteri at E9.5, RNA-seq of CD45^+^IL-18R^+^ cells purified from the uterine myometrium using a cell sorter at E13.5, examination of pups and placentas delivered by cesarean section at E18.5, and evaluation of fetal growth and neurological development after spontaneous delivery (Figure 4A).

**Figure 4.**
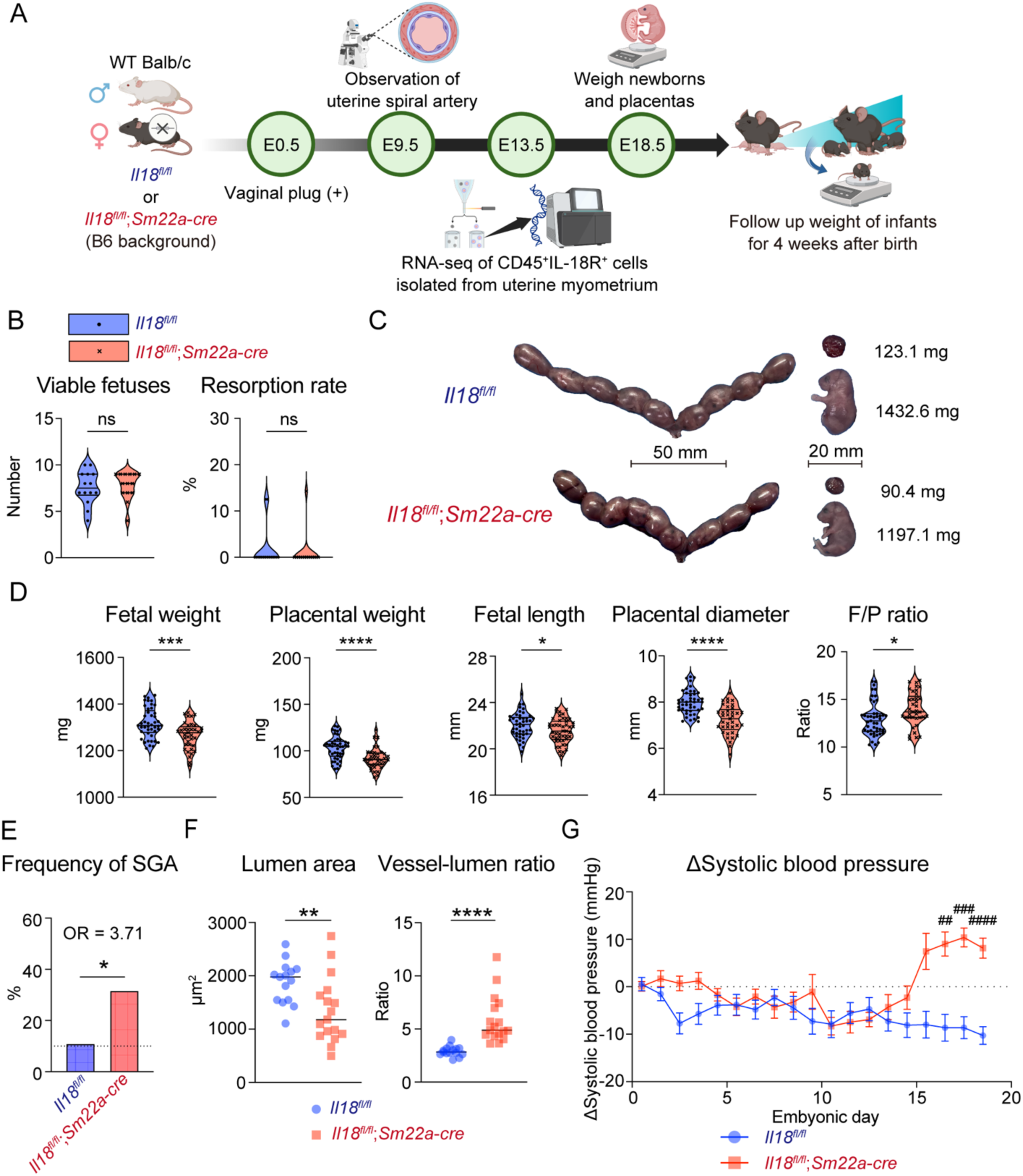
SMC-specific *Il18* KO mice exhibit a preeclampsia-FGR-like clinical course. (A) Schema of the experiments using *Il18^fl/fl^;Sm22a-cre* and *Il18^fl/fl^* mice is shown. Created with BioRender.com. (B) Violin plots representing the number of viable fetuses and the frequency of spontaneous embryo resorption at E13.5 or E18.5 are shown by group. (Mann–Whitney U test, each data point represents 1 dam, n = 12 per each group, each horizontal line represents median and the first and third quartile of samples). (C) Representative pictures of pregnant uteri, placentas, and newborns at E18.5 are shown with the weight of placentas and newborns by group. (D) Violin plots representing fetal weight, placental weight, fetal length, and placental diameter at E18.5 are shown by group. (Student’s *t*-test, each data point represents 1 newborn or placenta, 5 dams per each group, each horizontal line represents median and the first and third quartile of samples). (E) Frequency of small for gestational age (SGA) at E18.5 is shown by group. The SGA cutoff was defined as the 10th percentile of newborns from *Il18^f/f^* dams (Fisher’s exact test, OR = odds ratio). (F) Quantification of spiral artery remodeling with vessel lumen area and ratio between vessel wall area and vessel lumen area (Mann–Whitney U test, each data point represents the mean value of 15 measurements, 15–16 implantation sites from 5 pregnancies, each horizontal line represents the mean value of measurements). (G) The change in maternal systolic blood pressure relative to the premating baseline of each mouse is shown for each group, from E0.5 to E18.5. (Multiple Mann–Whitney U test, each data point represents the mean value of 11–13 dams per each group, each horizontal line represents ±standard error of mean) * *p* < 0.05, ** *p* < 0.01, *** *p* < 0.001, **** *p* < 0.0001 ## *q* < 0.01, ### *q* < 0.001, ### *q* < 0.0001 Related to Figure S4.

Although the litter size or spontaneous embryo resorption rates showed no differences between the two groups of dams (Figure 4B), the fetuses and placentas from *Il18^fl/fl^;Sm22a-cre* dams were significantly smaller and lighter (Figures 4C, D). Using the 10th percentile body weight of offspring born to *Il18^fl/fl^* dam as the reference, the odds ratio for small for gestational age (SGA) in *Il18^fl/fl^;Sm22a-cre* offspring increased by 3.71-fold (Figure 4E). Similarly, macrophage-specific *Il18* conditional KO mice (*Il18^fl/fl^;LysM-cre*), which were also generated using the Cre-loxP system, exhibited a similar tendency toward lower fetal and placental weights, emphasizing that IL-18 promotes fetal and placental growth (Figure S4F).

We next investigated uterine spiral artery remodeling, which is critical for maternal oxygen and nutrient supply contributing to placental development. In *Il18^fl/fl^;Sm22a-cre* dams, the spiral arteries exhibited a narrow lumen and thick vessel walls, indicating poor remodeling (Figures 4F, S4G), which is a typical finding in preeclampsia and FGR. In fact, compared with that in *Il18^fl/fl^* dams, *Il18^fl/fl^;Sm22a-cre* dams demonstrated a significant increase in systolic blood pressure in the late phase of pregnancy (Figure 4G). Conversely, measurements of maternal serum concentrations of soluble Fms-related receptor tyrosine kinase-1 (sFLT-1) and placental growth factor (PlGF) revealed results that appeared to contradict the classical “two-stage” theory of preeclampsia pathogenesis (Figure S4H). This theory postulates a first stage characterized by abnormal placentation and impaired uteroplacental perfusion, followed by a second stage involving decreased placental production of PlGF and increased production of sFLT-1, which neutralizes PlGF, ultimately worsening the disease and causing maternal multiorgan dysfunction ^33,34^. Unlike the typical preeclampsia-like phenotypes described thus far, *Il18^fl/fl^;Sm22a-cre* dam mice exhibited higher serum PlGF levels and unaltered sFLT-1 levels compared with *Il18^fl/fl^* mice, resulting in a lower sFLT-1/PlGF ratio (Figure S4H). This result may be attributed to the timing of blood sampling before the development of changes in the sFLT-1/PlGF ratio during the course of preeclampsia or a more prominent upregulation of PlGF production in response to excessive inflammation or hypoxia ^35^.

Observations of postnatal growth revealed that pups born smaller to *Il18^fl/fl^;Sm22a-cre* dams exhibited catch-up growth by 1 week of age. At 2–3 weeks of age, these pups even tended to weigh more than those born to *Il18^fl/fl^* dams (Figure S4I). This pattern is consistent with findings from a previous study of a preeclampsia mouse model expressing human sFLT-1 ^36^ and suggests that predictive adaptive responses to FGR contribute to metabolic abnormalities in the next generation. Moreover, neuromotor development appeared to be delayed in offspring born to *Il18^fl/fl^;Sm22a-cre* dams (Figure S4J).

Altogether, these findings indicate that IL-18 derived from SMCs plays a vital role in placental and fetal development by remodeling the maternal uterine spiral artery. Furthermore, *Il18^fl/fl^;Sm22a-cre* mice exhibited a preeclampsia/FGR-like clinical course compared with control mice.

### IL-18-signaling modulates uterine cytokine signaling pathways and NK cell function

To clarify the effects of SMC-derived IL-18 on the uterine immune milieu, we performed RNA-seq on CD45⁺IL-18R⁺ cells isolated using a cell sorter from the pregnant uterine myometrium of either *Il18^fl/fl^;Sm22a-cre* or *Il18^fl/fl^* mice (Figures 4A, S5A). Although the CD45⁺IL-18R⁺ cell population contained the highest proportion of NK cells (Figure S5B), this finding is consistent with previous results showing that despite NK cells constituting a relatively small population in the uterine myometrium, they exhibited an extremely high IL-18R positivity rate (Figures 1B, 1D, 1E). We normalized the RNA-seq raw count data using DEseq2 (Figure S5C) and found that the two groups clearly separated along the second principal component (Figure 5A). Genes with a *q*-value <0.1 and an absolute fold change >1.5 were used to identify genes that were significantly upregulated or downregulated in *Il18^fl/fl^;Sm22a-cre* mice compared with that in *Il18^fl/fl^* mice (Figures 5B, 5C, S5C; Tables S1A, S1B).

**Figure 5.**
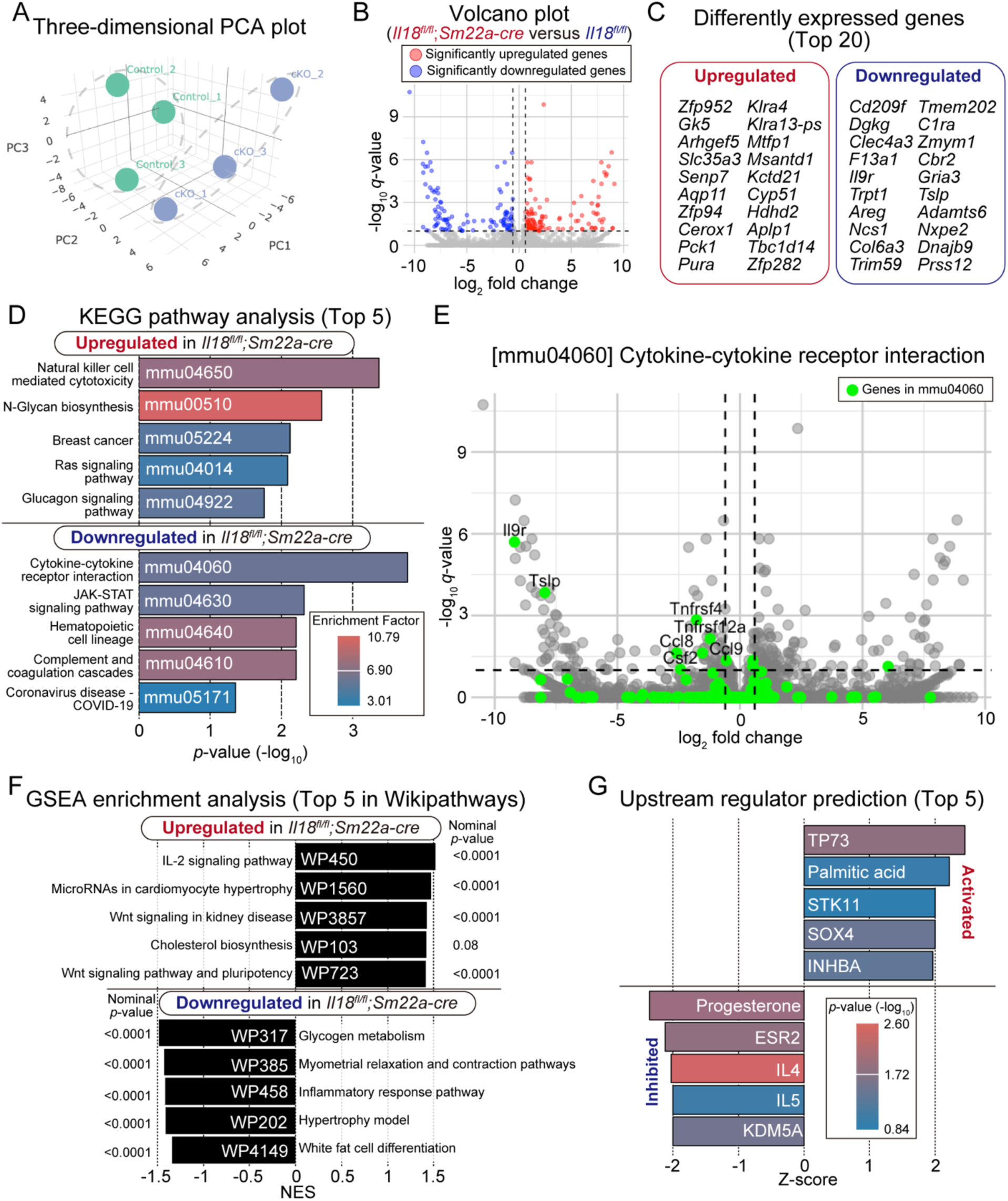
RNA-seq reveals the impact of IL-18-signaling on the immune milieu. (A) Principal component analysis scores three-dimensional plots of RNA-seq of samples (n = 3 per group). (B) Volcano plots depicting the results of RNA-seq analysis, with highlighted plots representing significantly upregulated or downregulated genes. (C) The top 20 significantly upregulated or downregulated genes sorted by *q*-value are shown. Noncoding RNAs are excluded from the list (Related to Table S1). (D) The top 5 upregulated or downregulated pathways identified by KEGG pathway analysis are shown sorted by *p*-value. The bar plot color indicates the heatmap of the enrichment factor (Related to Table S2). (E) Volcano plots depicting the results of RNA-seq analysis, with highlighted plots representing the mmu04060 gene set. Significantly downregulated genes in the gene set were labeled with the gene symbol. (F) Top 5 upregulated or downregulated pathways identified by gene set enrichment analysis (GSEA) using Wikipathways are shown sorted by normalized enrichment score. The nominal *p*-value for each pathway is shown together (Related to Tables S3, Table S4). (G) Top 5 activated or inhibited regulators predicted by IPA are shown sorted by Z-score. The bar plot color indicates the heatmap of *p*-value (Related to Table S5). Related to Figure S5.

KEGG pathway analysis of downregulated genes revealed that gene sets such as “Cytokine–cytokine receptor interaction [mmu04060]” and “JAK–STAT signaling pathway [mmu04630]” were among the most enriched (Figure 5D; Table S2B). Within the [mmu04060] gene set, a variety of genes were downregulated, including *Csf2*, which encodes GM-CSF produced by CD4⁺ T cells under Th1 condition, and Tslp, which encodes thymic stromal lymphopoietin that facilitates the Th2 differentiation of CD4⁺ T cells (Figure 5E). These findings suggest the involvement of IL-18-signaling in the uterine immune environment in the regulation of both type 1 and type 2 immune responses.

KEGG pathway analysis of upregulated genes revealed that “Natural killer cell mediated cytotoxicity [mmu04650]” was most highly enriched (Figure 5D; Table S2A). Gene set enrichment analysis (GSEA) using the most similar Gene Ontology annotation, “Natural killer cell mediated immunity [GO:0002228]”, also revealed an upregulation of genes encoding killer cell lectin-like receptors—such as *Klrd1* and *Klrc2*— which act as activating receptors on NK cells (Figures S5D, S5E; Table S4A). In reproductive immunology, it is well established that NK cells in the uterus exhibit a unique differentiation pattern, with uterine NK cells supporting pregnancy through the production of factors such as GM-CSF and VEGF and conventional NK cells releasing cytotoxic granules containing perforin and granzyme that can damage the placenta ^37^. In fact, in *Il18^fl/fl^;Sm22a-cre* mice, the expression of *Csf3* significantly reduced, whereas that of genes encoding granzyme and perforin tended to increase (Figure S5E). These findings suggest that IL-18- signaling plays a critical role in modulating NK cell function, warranting further research on NK cell subtypes and their functions. Moreover, “N-Glycan biosynthesis [mmu00510]” pathway was also highly enriched in *Il18^fl/fl^;Sm22a-cre* mice (Figure 5D; Table S2A). The process of N-glycosylation is essential for proper placental development by trophoblast cells and for the maintenance of maternal–fetal immune tolerance; however, excessive or aberrant patterns of N-glycosylation have been implicated in the pathogenesis of preeclampsia ^38–40^.

Using GSEA with the open-access database Wikipathways, which has abundant pregnancy-related gene sets, the “IL-2 signaling pathway [WP450]” was found to be highly ranked (Figure 5F; Table S3A). Increased IL-2 production has been reported in preeclampsia and recurrent miscarriage, and our results remarkably revealed an upregulation of MAPK-related genes within this gene set (Figure S5F; Table S4B). Conversely, among the downregulated gene sets, “Myometrial relaxation and contraction pathways [WP385]” ranked highly (Figure 5F, Table S3B). The observed downregulation of genes associated with both contraction and relaxation of the uterine myometrium suggests that IL-18 derived from uterine SMCs also affects the contraction and relaxation of the myometrium (Figure S5G; Table S4C).

Finally, upstream regulator prediction using Ingenuity Pathway Analysis (IPA) identified progesterone (P4) as one of the top inhibited upstream factors (Figure 5G; Tables S5A, S5B). Although this may be due to decreased P4 production by the placenta resulting from placental insufficiency, P4-PR signaling is known to regulate the functions of numerous uterine immune cells ^41^, raising the possibility that P4 acts as a homeostatic inducer of IL-18-signaling.

To summarize, these findings suggest that deficient IL-18-signaling in the uterus suppresses various cytokine signaling pathways and, in particular, promotes the cytotoxicity of NK cells.

### IL-18 promotes placental angiogenesis *in vitro*

According to the database of The Human Protein Atlas (HPA) ^42^, the expression of *IL18R1*, which encodes the human IL-18R was highest in the placenta among all systemic tissues (Figure S6A). Considering that multiple reports suggest that IL-18 promotes angiogenesis either indirectly or directly ^43,44^, we hypothesized that IL-18 supports placental growth by promoting placental angiogenesis.

We conducted a tube formation assay using mouse primary placental microvascular endothelial cells (PVECs) treated with vehicle, recombinant mouse IL-18 (rIL-18), or recombinant mouse VEGF (rVEGF) (Figure 6A). In fact, PVECs exhibited a moderate IL-18R positivity compared with other uterine immune cells (Figures 1E, S6B). Consequently, the treatment with rIL-18 promoted angiogenesis in PVECs to a similar degree as that in the positive control treated with rVEGF (Figure 6B). The images were quantified for the extent of angiogenesis using the Angiogenesis Analyzer software ^45^ in ImageJ ^46^. Compared with that in vehicle, rIL-18 treatment significantly promoted angiogenesis in a dose-dependent manner (Figures 6C, S6C).

**Figure 6.**
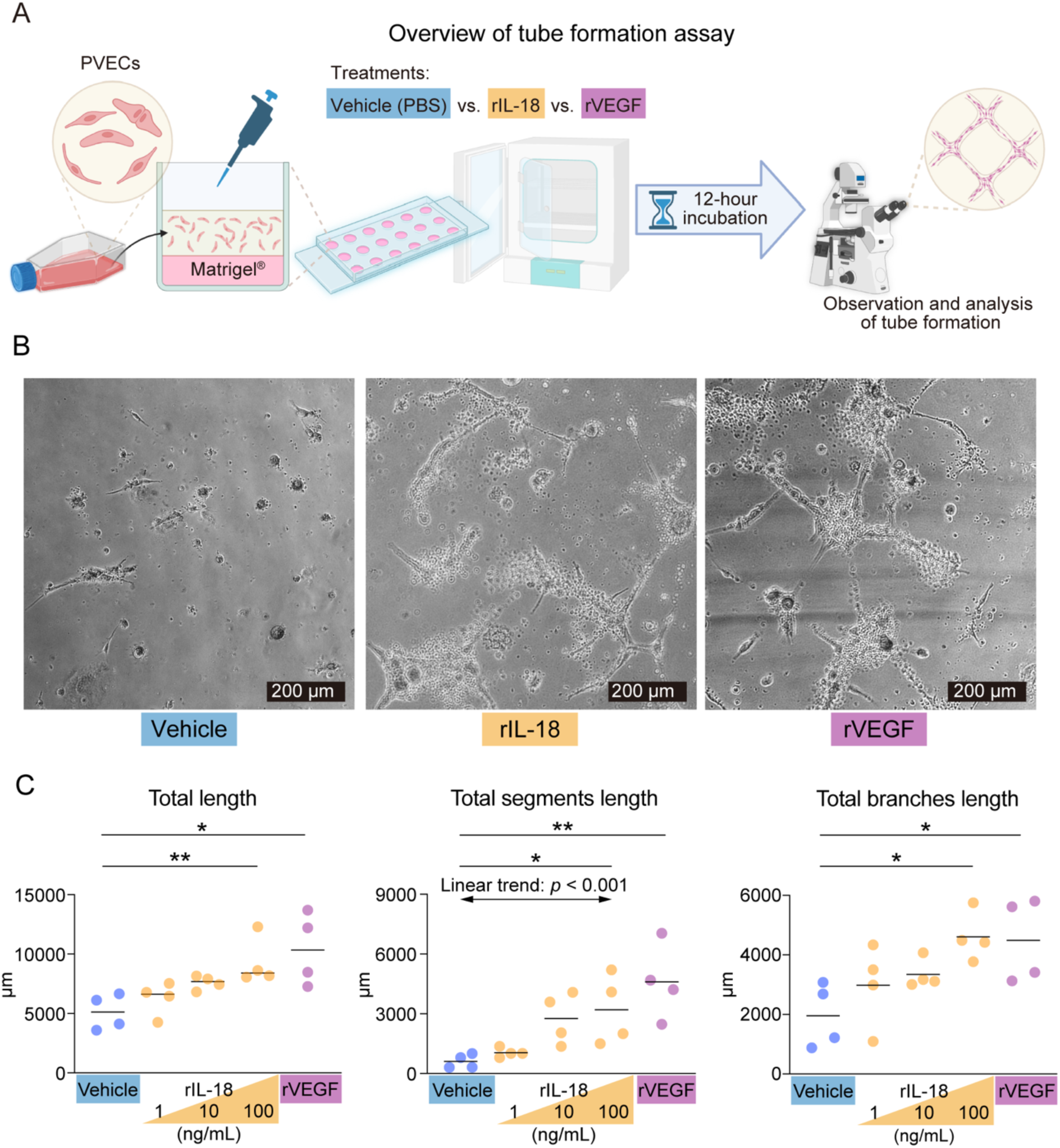
IL-18 promotes placental angiogenesis *in vitro*. (A) Schematic illustrating the tube formation assay using mouse primary placental microvascular endothelial cells (PVECs). Created with BioRender.com. (B) Representative images of tube formations observed by phase contrast microscopy are shown for each treatment. (C) Images obtained by the tube formation assay were quantified using the Angiogenesis Analyzer for ImageJ and plotted for each indicated parameter by each treatment (one-way analysis of variance, linear trends from vehicle to 100 ng/mL rIL-18 treatment were also analyzed, each data point represents 1 sample, n = 4 per each group, each horizontal line represents the mean value of samples). * *p* < 0.05, ** *p* < 0.01 Related to Figure S6.

In summary, these findings indicate that IL-18 supports placental development not only through maternal tissue remodeling but also by promoting placental angiogenesis.

## Discussion

This study clarified that IL-18, which is abundantly produced in the pregnant mouse uterine myometrium, controls the immune profile of the uterine immune milieu and plays a vital role in fetal development through uterine vascular remodeling and placental formation. Remarkably, *Il18^fl/fl^;Sm22a-cre* dam mice exhibited the following preeclampsia-like phenotypes: restriction of fetoplacental growth, poor remodeling of the maternal uterine spiral artery, elevated maternal blood pressure, and delayed neurodevelopment and excessive weight gain in offsprings. Both *in vivo* and *in vitro* experiments demonstrated that IL-18 promotes constitutive IFN-γ production by T cells and NK cells, which express high levels of IL-18R, and is involved in maintaining the M1 macrophage profile and IL-12 production. These data suggest that IL-18 supports the type 1 immune response required for the maternal tissue remodeling.

Our recent study using a low-dose lipopolysaccharide (LPS)-induced abortion model to investigate the miscarriage-preventive effect of IL-18 has found that miscarriages induced at a high rate by low-dose LPS, and IL-18 nAb were rescued by combination therapy with recombinant IFN-γ and recombinant IL-4 (manuscript under review). In the present study, our *in vitro* experiments also demonstrated that administration of IL-18 nAb suppressed the production of both IFN-γ and IL-4 by uterine myometrial immune cells, and similarly, both type 1 and type 2 immune response signaling were downregulated in the RNA-seq analysis of *Il18^fl/fl^;Sm22a-cre* mice. Although discussions of reproductive immunology often focus on a unidirectional type 1 or type 2 immune response, our results suggest that both type 1 and type 2 immune responses are concurrently driven by cytokines such as IL-18 that exert dual effects.

IL-18 is often considered a proinflammatory cytokine harmful to pregnancy, and in fact, elevated IL-18 levels have been reported in cases of recurrent miscarriage and preterm birth ^47^. It remains unclear whether IL-18 is merely a byproduct of an excessive inflammatory response resulting in adverse pregnancy outcomes or whether an excessive IL-18 level itself causes such adverse outcomes. Meanwhile, other studies have indicated that dysbiosis of the gut microbiota suppresses IL-18 production in hepatic Kupffer cells, thereby hindering NK cell maturation ^48^, and that IL-18-signaling is inhibited in uterine adenomyosis lesions ^49^. Gut dysbiosis ^50–52^ and uterine adenomyosis ^53,54^ have been clinically reported to correlate strongly with preeclampsia. Although it is unlikely that excessive IL-18 levels would exert a positive effect on pregnancy, inhibition of homeostatic IL-18-signaling suggests a negative impact on the immune environment and pregnancy, which is consistent with our experimental results.

In conclusion, we demonstrated that IL-18 produced by pregnant uterine SMCs regulates the uterine immune milieu and thereby promotes fetoplacental growth. Moreover, examining maternal immune environments and pregnancy outcomes with a focus on IL-18-signaling may enable earlier prediction of risks such as preeclampsia and miscarriage/preterm labor and provide novel preconception care strategies and therapeutic interventions.

## Limitations of the study

A limitation of our study is that the detailed mechanisms underlying IL-18 production in uterine SMCs remain incompletely understood. Although we detected the activation of inflammasome-related proteins in the uterine SMCs, inflammasome formation generally requires robust inflammatory stimuli like LPS which is unlikely under normal pregnancy conditions. We performed IPA using our RNA-seq data, which predicted suppression of progesterone and ESR2 (estrogen receptor 2) in *Il18^fl/fl^;Sm22a-cre* mice. These findings suggest that pregnancy-specific hormonal exposure and signaling may contribute to the homeostatic IL-18 production. Moreover, inflammasome-independent mechanisms may also be involved. ADAM33, a zinc-dependent metalloprotease capable of cleaving pro-IL-18 into its mature form independently ^55^, has been reported to be highly expressed in the uterus compared to other human tissues^42^ (https://www.proteinatlas.org/ENSG00000149451-ADAM33/tissue). These suggests that IL-18 production might involve multiple potential pathways.

Another further limitation is the complexity of IL-18-signaling itself, including unique regulatory molecules such as IL-18-binding proteins and IL-18 receptor accessory proteins. These were not explored in detail but require further investigation to completely clarify the role of IL-18 in the reproductive immune milieu. Finally, although our results emphasize a significant effect of IL-18 on uterine NK cells, our study did not investigate these NK cell subsets in depth. Future research focusing on IL-18-mediated NK cell functional diversity would probably provide important insights into uterine immune regulation.

## Data availability

RNA-seq raw data have been deposited in the National Center for Biotechnology Information (NCBI) Gene Expression Omnibus (GEO); the accession number is GSE293908. These data are publicly available.

The data that support the findings of this study are available from the lead contact, R.M., upon reasonable request.

## Acknowledgements

We sincerely appreciate Ms. Masumi Shimizu and Ms. Miyuki Takatori for their technical support and all the staff of the Department of Microbiology and Immunology at Nippon Medical School Graduate School. Scientific images were provided by BioRender (https://www.biorender.com/). This research was funded by the Japan Society for the Promotion of Science (JSPS) KAKENHI (Grant number 20K09679 and 24K12637 to Y.N.; 16H05178 and 22K07140 to R.M.), the Nippon Medical School Grant-in-Aid for Medical Research, the Naito Foundation, the Daiichi Sankyo Foundation of Life Science, the Uehara Memorial Foundation, the Takeda Science Foundation, and the Terumo Life Science Foundation.

## Author contributions

H.I., Y.N., and R.M. designed the project. H.I., Y.N., Y.H., and E.K. conducted the experiments. H.I., Y.H., E.K., and R.F. generated the conditional knockout mice. H.I., Y.N., S.S., and R.M. contributed to data interpretation, while S.S. and R.M. offered scientific guidance and supervised the entire study. All authors took part in revising the manuscript, final approval for publication, and accepted responsibility for the integrity of the work.

## Declaration of interests

The authors declare that they have no known competing financial interests or personal relationships that could have appeared to influence the work reported in this paper.

## Materials and Methods

### Key resources table

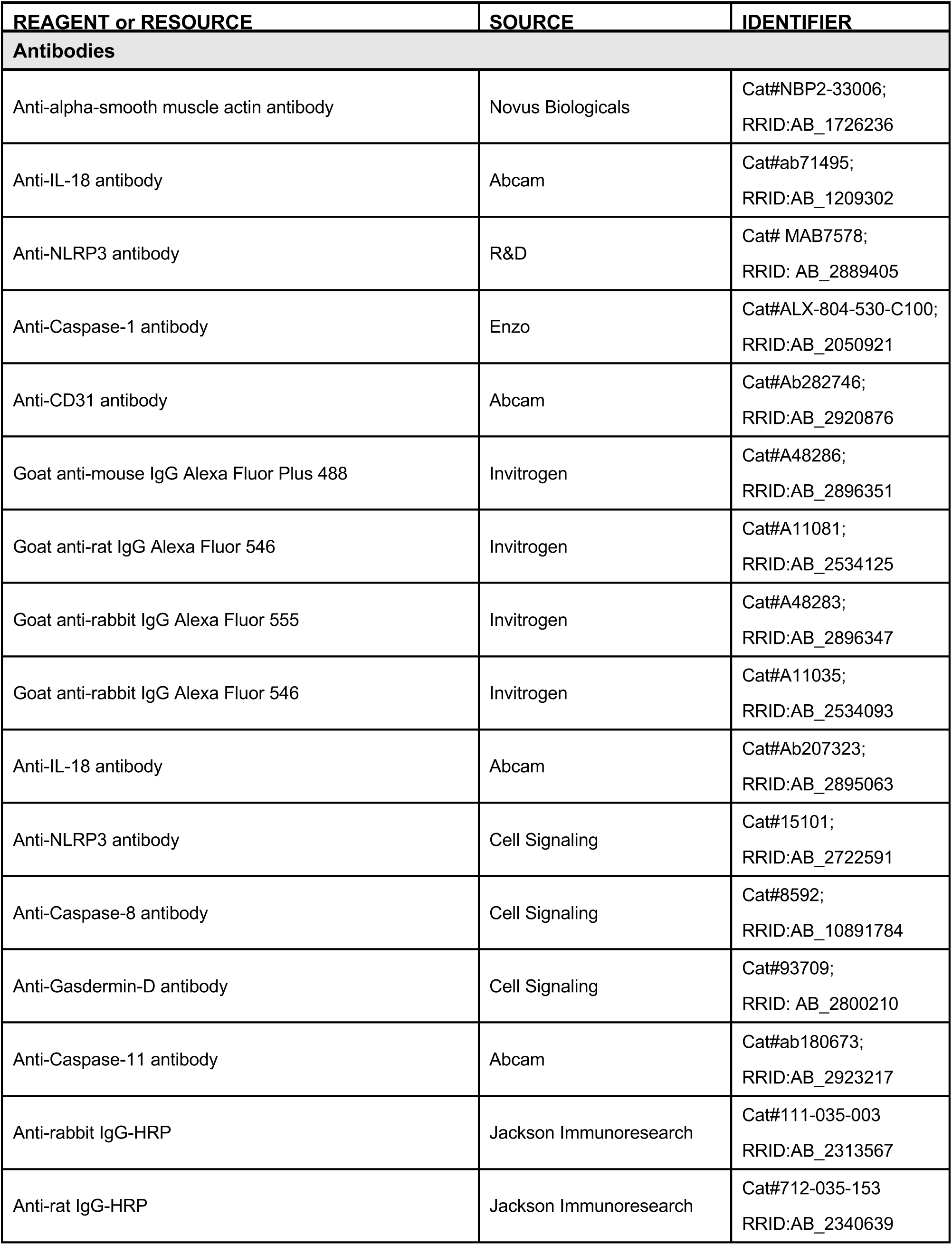
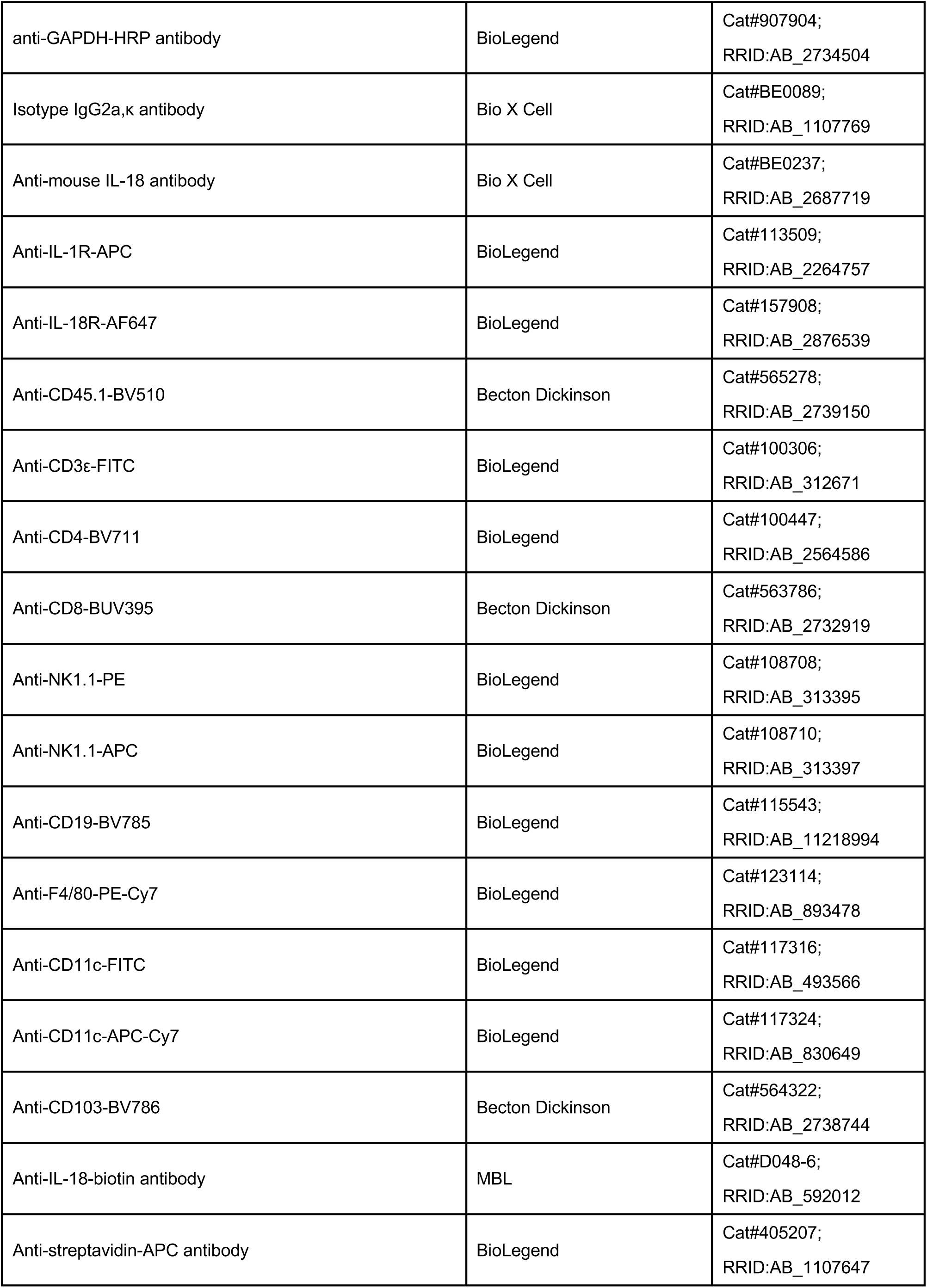
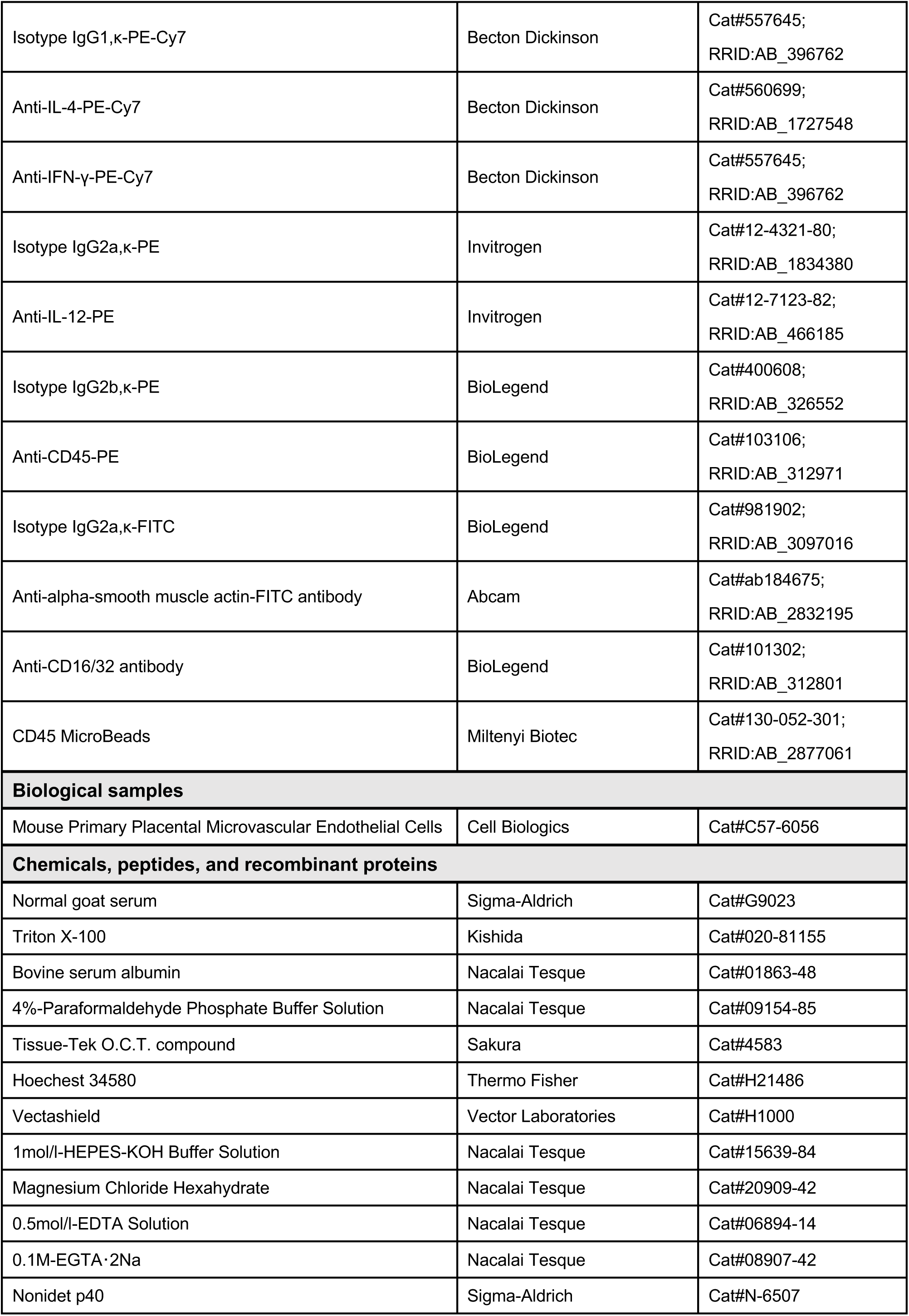
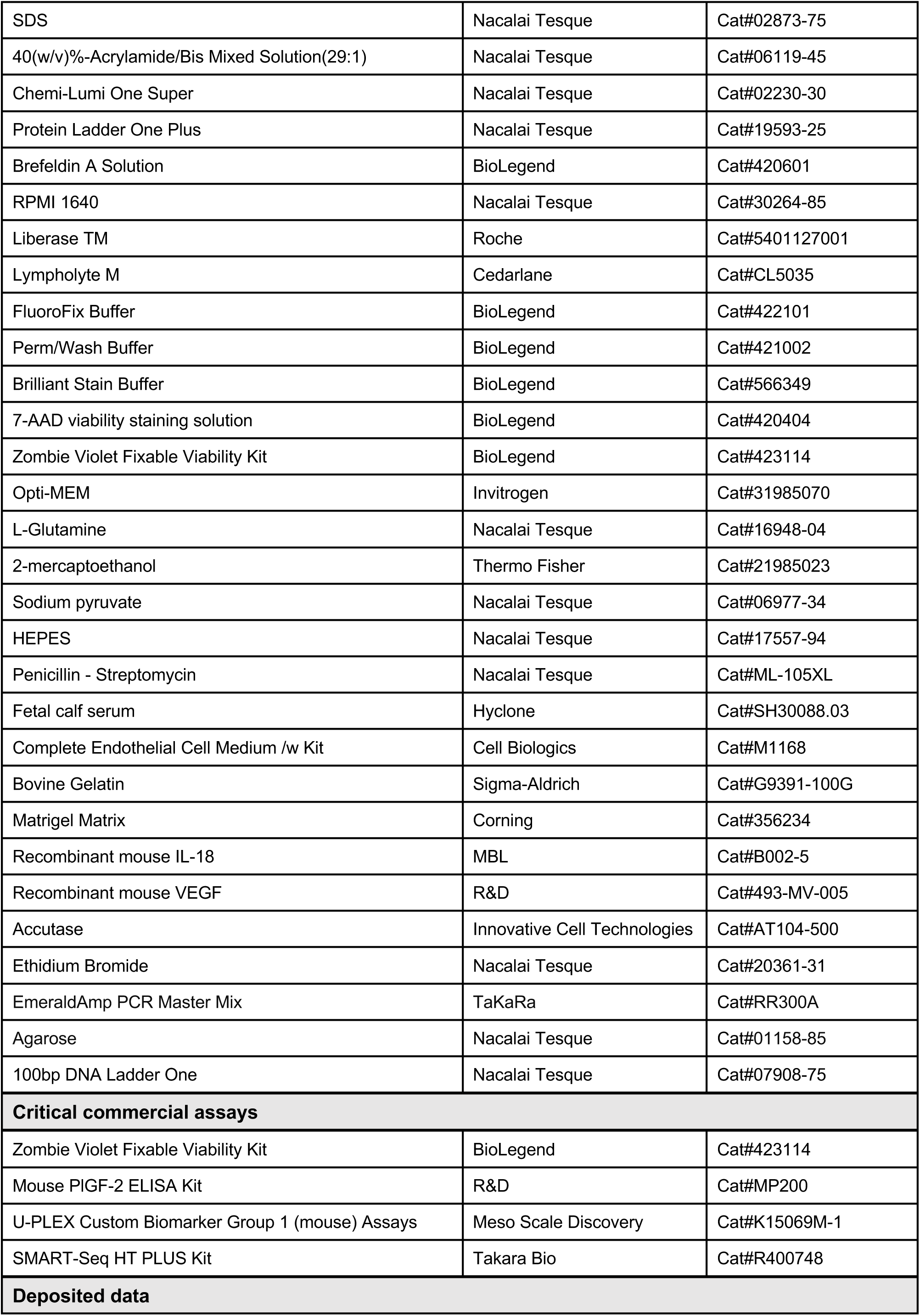
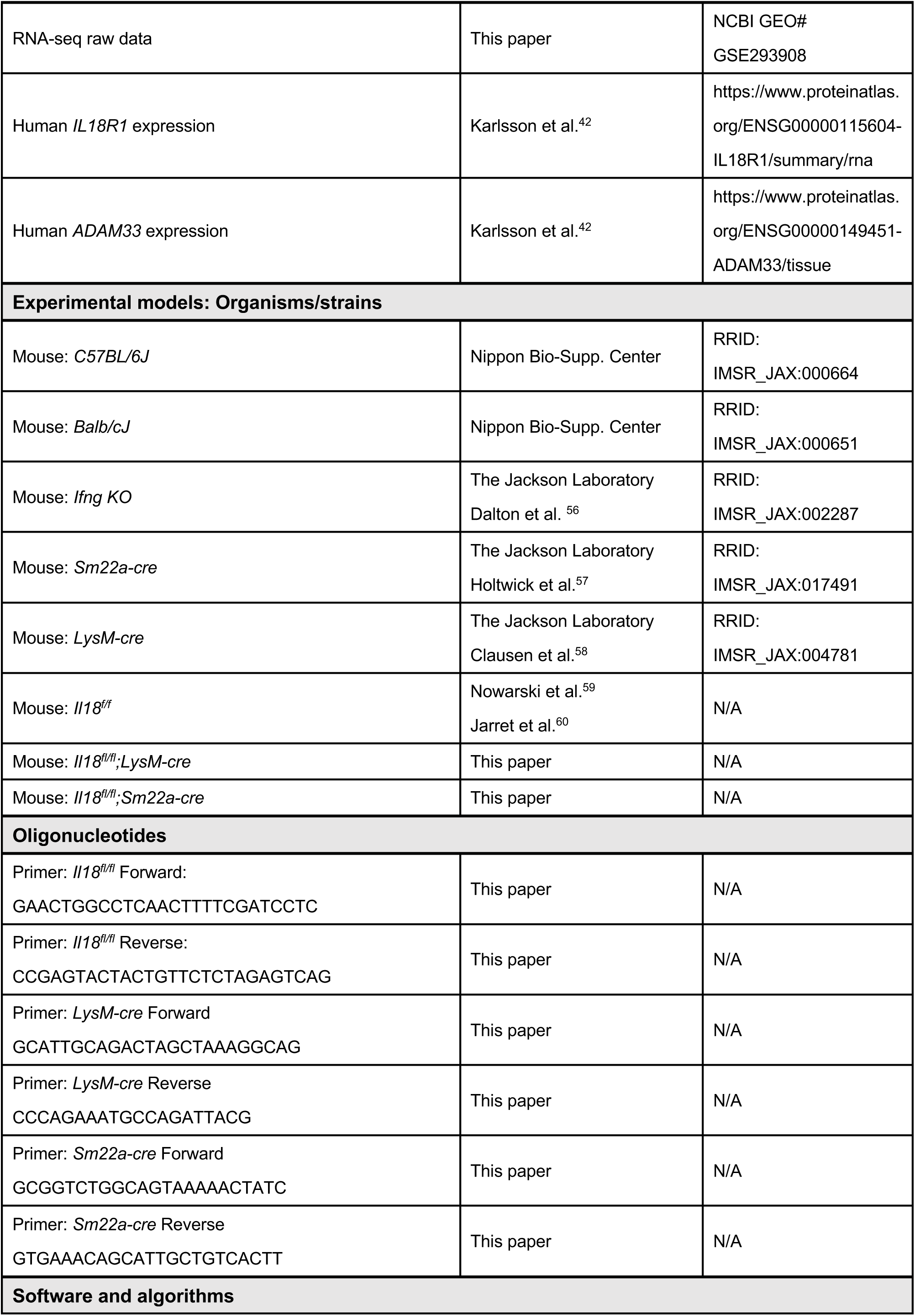
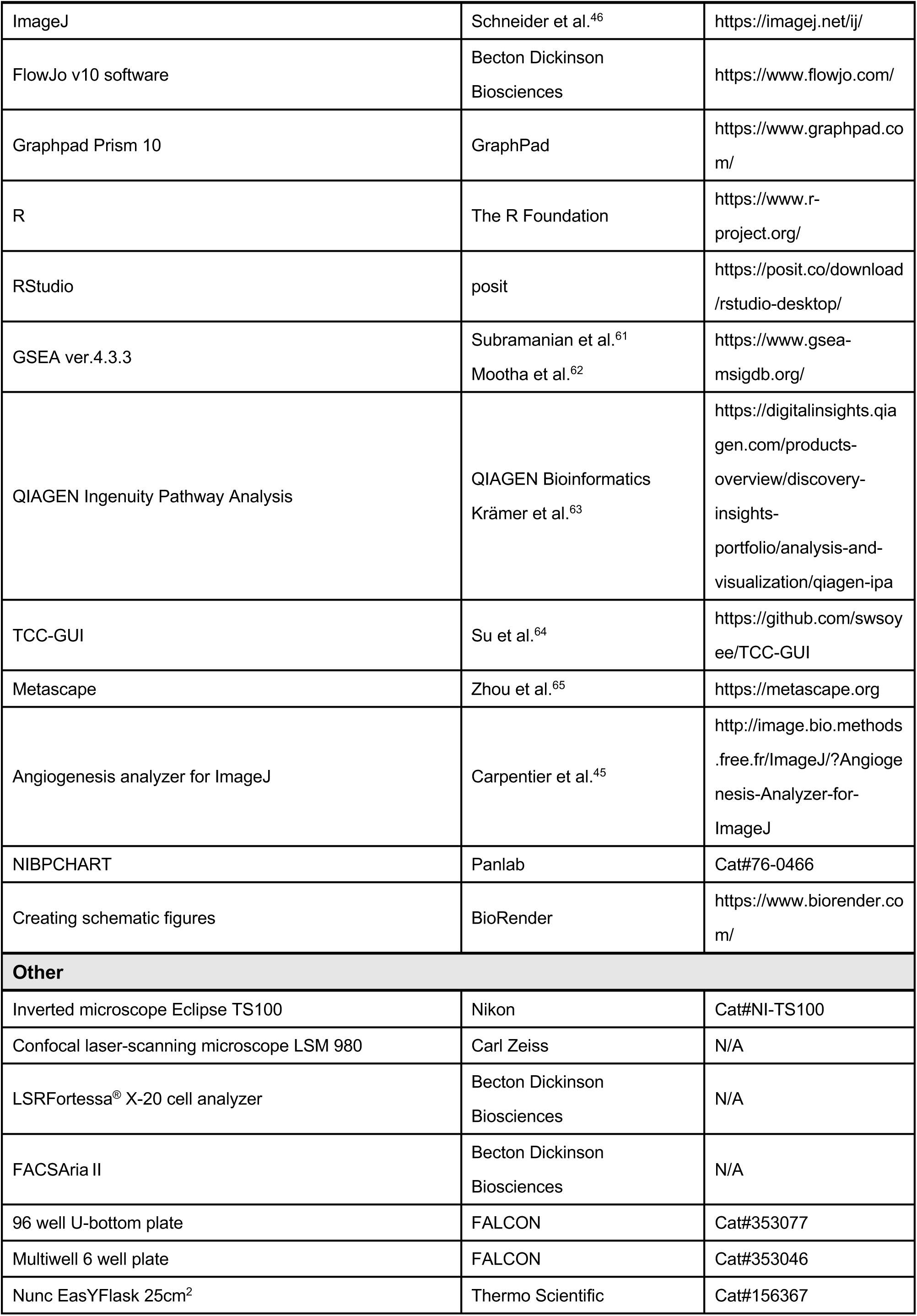
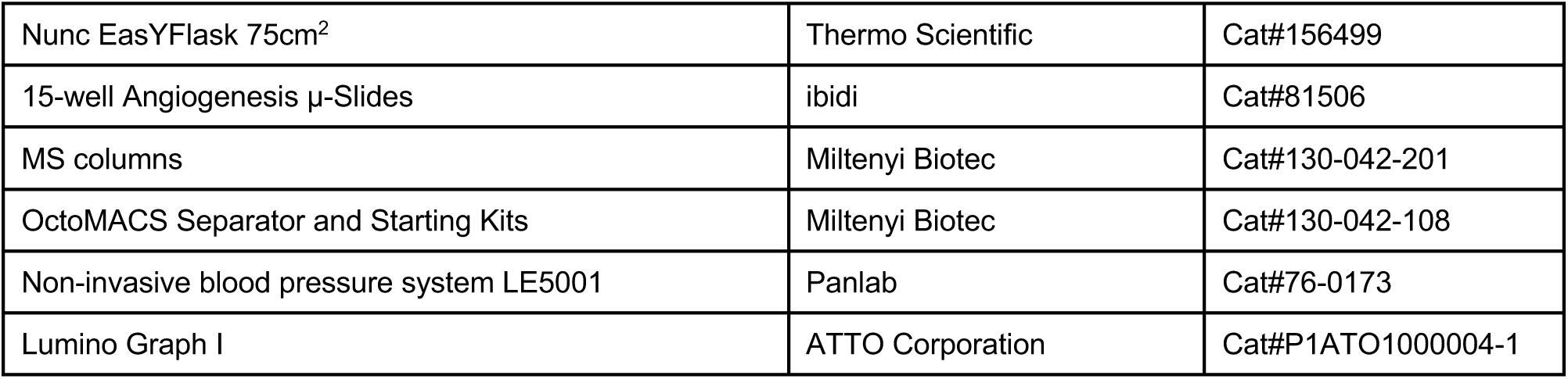

### Animals and breeding

C57BL/6J (WT) and BALB/cJ (WT) mice were obtained from Nippon Bio-Supp. Center Co., Ltd. (Tokyo, Japan). *Ifng* KO ^56^ (strain: IMSR_JAX:002287), *Sm22a-cre* ^57^ (strain: IMSR_JAX:017491) and *LysM-cre* ^58^ (strain: IMSR_JAX:004781) mice were on a C57BL/6 background and were obtained from the Jackson Laboratory (ME, USA). *Il18^fl/fl^* mice on a C57BL/6J background ^59,60^ were bred with *Sm22a-cre* mice to generate *Il18^fl/fl^;Sm22a-cre* mice as SMC-specific *Il18* conditional knockout mice. *Il18^fl/fl^;LysM-cre* mice were similarly generated as macrophage-specific *Il18* mice conditional knockout mice by crossing *Il18^fl/fl^* and *LysM-cre* mice. All mice were bred and maintained in a specific pathogen free area of the experimental medicine animal room at Nippon Medical School (Tokyo, Japan). A circadian cycle (light: dark = 12: 12 h) was used for all mice. Female mice aged 8–12 weeks were mated with male mice in the late afternoon. All female mice on a C57BL/6 background were allogeneically mated with male BALB/c mice. Vaginal plugs were checked daily between 8 AM and 9 AM, and the appearance of the vaginal plug indicated E0.5. All animal experiments were conducted according to the Guidelines for the Care and Use of Laboratory Animals issued by the National Institutes of Health (NIH, Bethesda, MD, USA), and all protocols were approved in advance by the Animal Experimental Ethical Review Committee of Nippon Medical School (approval No. 2022-010). The study methods were performed and reported in agreement with the ARRIVE guidelines.

### Genotyping of conditional knockout mice

To confirm the genotype of theconditional knockout mice, genomic DNA was extracted from the tail tip using an alkaline lysis method. The reaction was neutralized using Tris–HCl, and the supernatant was used as a template for polymerase chain reaction (PCR). Amplified PCR fragments were separated by size on a 1% agarose gel and visualized using ethidium bromide staining. The primer sequences used for genotyping are listed in the key resources table.

### Sample collection and preparation

For cell analysis, 4 h before euthanasia, brefeldin A (BioLegend, # 420601) was administered via intraperitoneal injection. On the designated gestational day for each experiment, the mice were euthanized by cervical dislocation. An abdominal incision was made using ophthalmic scissors, and the pregnant uterus was excised, excluding the cervix. The uterine body was incised, and the myometrium, decidua, placenta, and fetuses were carefully separated.

Before cell analysis and *in vitro* experiments, the tissues were cut into small pieces and incubated with RPMI1640 (Nacalai Tesque, #30263-95) and 5 mg/mL LiberaseTM (Roche, #05401127001) at 37°C for 1 h. Next the tissues were filtered through a nylon mesh, and the viable cells were isolated from dead cells and debris using Lympholyte-M (Cedarlane, #CL5035).

Before western blotting, after flash-freezing each tissue in liquid nitrogen and crushing it, the samples were lysed in a sample buffer containing 50 mM HEPES/KOH (Nacalai Tesque, #15639-84), 250 mM NaCl, 1.5 M MgCl₂ (Nacalai Tesque, #20909-42), 1 mM EDTA (Nacalai Tesque, #06894-14), 1 mM EGTA (Nacalai Tesque, #08907-42), and 0.5% Nonidet P-40 (Sigma-Aldrich, #N-6507).

### Western blotting analysis

Whole-cell lysates were subjected to SDS-PAGE under reducing conditions and transferred onto polyvinylidene fluoride membranes. After blocking, the membranes were incubated with a monoclonal antibody. Next, they were treated with horseradish peroxidase–conjugated secondary antibodies, and then visualized using Chemi-Lumi One Super (Nacalai Tesque, #02230-30) together with LiminoGraph Ⅰ (ATTO Corporation, #P1ATO1000004-1), according to the manufacturers’ instructions. Antibodies used in the western blotting are listed in the key resources table.

### Flow cytometric analysis and sorting

Cells were washed with flow cytometry washing buffer containing 3% fetal bovine serum and then the cells were stained with a mixture of fluorescent-conjugated antibody and Brilliant Stain Buffer (BD, #563794) for extracellular staining. After fixation and permeabilization, the cells were intracellularly stained with a fluorescent conjugated antibody. For indirect staining, the cells were incubated with a secondary antibody, after incubating with the first antibody. For negative selection of dead cells, 7-aminoactinomycin D (BioLegend, #420404) was used for extracellular staining and Zombie Violet Fixable Viability Kit (BioLegend, #423114) was used for intracellular staining. Gating was performed by removing debris, dead cells, and doublets, after which immune and nonimmune cells were separated based on CD45 negativity or positivity. Each immune cell population was defined as follows: T cells (CD3^+^), CD4^+^ T cells (CD3^+^CD4^+^), CD8^+^ T cells (CD3^+^CD8^+^), B cells (CD3^-^CD19^+^), NK cells (CD3^-^NK1.1^+^), macrophages (F4/80^+^), M1 macrophages (F4/80^+^CD11c^+^), M2 macrophages (F4/80^+^CD11c^-^), DCs (F4/80^-^CD11c^+^), DC1 (F4/80^-^ CD11c^+^CD103^+^), and DC2 (F4/80^-^CD11c^+^CD103^-^). The stained cells were analyzed using a BD LSR Fortessa X-20 flow cytometer (Becton Dickinson Biosciences) and FlowJo software v10.10 (Becton Dickinson) and sorted using a BD ARIA II (Becton Dickinson Immunochemical System). Antibodies used in the flowcytometry are listed in the key resources table.

### *In vitro* assay using uterine cells isolated using MACS

After incubation with Fc-blocker (BioLegend, #101302), CD45-positive cells were purified magnetically using anti-CD45 Micro Beads (positive selection; Miltenyi Biotec, #130-052-301) through an MS column (Miltenyi Biotec, #130-042-201) and then suspended in RPMI-1640-based culture medium supplemented with 0.1 mM nonessential amino acids (Invitrogen, #31985070), 2 mM L-glutamine (Nacalai Tesque, #16948-04), 50 mM 2-mercaptoethanol (Thermo Fisher, #21985023), 1 mM sodium pyruvate (Nacalai Tesque, #06977-34), 1 mM HEPES (Nacalai Tesque, #17557-94), 100 U/mL penicillin-streptomycin (Nacalai Tesque, #ML-105XL), and 10% heat-inactivated fetal calf serum (Hyclone, #SH30088.03).

The cell suspension was dispensed into 96-well U-bottom multiwell plates at a concentration of 10^5^ cells per well and incubated with IL-18 nAb (Bio X Cell, #BE0237) or its isotype IgG (Bio X Cell, BE0089) under standard culture condition (37°C in 21% O2 and 5% CO2) for 12 h. The cultured cells were subjected to flow cytometry, and the supernatant was subjected to ELISA performed by a commercial service (St. Lukes SRL, Tokyo, Japan). Cytokine concentrations were quantified using U-PLEX Custom Biomarker Group 1 (mouse) Assay kit (MSD, Cat#k15069m).

### Immunofluorescence with two-dimensional Imaging

Pregnant mice were euthanized, and their uteri were extracted. The isolated implantation sites were embedded with Tissue-Tek O.C.T. compound (Sakura, Cat#4583), were frozen at –80°C and were cryosectioned at 7-µm thickness. Two or three implantation sites per dam were sampled, and two or three sections per implantation site were analyzed. The sections were fixed with 4% paraformaldehyde phosphate (Nacalai Tesque, Cat#09154-85) and were transparentized using 0.1% Triton X-100 (Kishida, #020-81155) before staining with the primary antibody. Then, the sections were blocked in phosphate buffered saline (PBS) containing 3% purified serum matching to the secondary antibody and stained with primary antibody overnight at 4°C. Negative control samples were blocked without the primary antibody. Next, the samples were incubated with the secondary antibody specifically recognizing the origin of the primary antibody for 30 min at room temperature. Nuclei were visualized using Hoechst 34580 (Thermo Fisher, Cat#H21486), and the sections were mounted using Vectashield (Vector Laboratories, #H1000). Imaging was performed using LSM980 (Carl Zeiss). Antibodies used in the immunofluorescent staining are listed in the key resources table.

### ELISA

Serum cytokine concentrations were measured using the ELISA kits according to the manufacturer’s instructions. ELISA kits are listed in the key resources table.

### Tube formation assay with placental vascular endothelial cells (PVEC)

Primary placental microvascular endothelial cells (CellBiologic, #C57-6056) were grown in gelatin-coated flasks with complete growth medium (CellBiologic, #M1168) under standard culture condition. For tube formation assay, 15-well angiogenesis µ-Slides (ibidi, # 81506) were previously coated with 10 µL Matrigel Matrix (Corning, # 356234). PVECs were applicated at a concentration of 10^4^ cells per well and was treated with vehicle (PBS), recombinant mouse IL-18 (MBL, #B002-5), or recombinant mouse VEGF (R&D, #493- MV-005). Photographs were taken at 12 h using a phase-contrast microscope. Quantification of tubular network structures was conducted using the Angiogenesis Analyzer software ^45^ (http://image.bio.methods.free.fr/ImageJ/?Angiogenesis-Analyzer-for-ImageJ) in ImageJ ^46^ (https://imagej.net/ij/).

### Spiral artery remodeling

Remodeling of spiral arteries was evaluated using methods that are described in detail in the literature ^21,66^. Briefly, E9.5 dams were euthanized, and the uterine artery was ligated at its distal and proximal ends. The entire uterus was extracted and fixed in 10% formalin and then dehydrated in 70% ethanol. The isolated implant sites were paraffin-embedded and sectioned at 7-μm thickness. Then, 49-μm-spaced sections were subjected to H&E staining, and for each implant site, three sections near the midsagittal point were analyzed. A total of 15 measurements of the decidual vessels were averaged to obtain the mean value for each implant site. The ImageJ software ^46^ was used to draw the outer wall and the vessel lumen to determine the wall area and the lumen area, respectively. The wall thickness was calculated by subtracting the lumen area from the wall, and the wall-to-lumen ratio was calculated by dividing the wall area by the lumen area. Three or four implant sites were analyzed for each dam.

### Gene expression analysis by RNA-seq

CD45⁺IL-18R⁺ cells isolated using BD ARIA Ⅱ (BD Biosciences) from pregnant uterine myometrium of either *Il18^fl/fl^;Sm22a-cre* or *Il18^fl/fl^* mice were stored at −80°C in a lysis buffer. Total RNA extraction, cDNA synthesis, and library generation were performed using the SMART-Seq HT PLUS Kit (Takara Bio, #R400748) according to the manufacturer’s instructions. Samples with a total RNA content of at least 1 ng and RNA fragments of at least 300 nucleotides were considered eligible. Pooled libraries were sequenced on an Illumina NovaSeq 6000 (Illumina) platform. The process from RNA extraction to gene expression analysis was performed by a commercial service (Rhelixa, Tokyo, Japan).

On the TCC-GUI platform ^64^ (https://github.com/swsoyee/TCC-GUI), raw read counts of 55,488 genes were standardized using DESeq2, low-count genes were excluded, and 24,061 genes were defined as eligible. GSEA ver.4.3.3 ^61,62^ (UC Sandiego and Broad Institute, https://www.gsea-msigdb.org/) was performed on all normalized counts. Genes with a *q*-value < 0.1 and a fold change >1.5 (127 genes) or <−1.5 (111 genes) were subjected to KEGG pathway analysis via the Metascape ^65^ (https://metascape.org) interface and QIAGEN IPA ^63^ (QIAGEN Bioinformatics, https://www.qiagenbioinformatics.com/products/ingenuity-pathway-analysis).

### Blood pressure measurement

Mouse blood pressure was measured using a noninvasive tail-cuff device LE5001 (Panlab, Cat#76-0173) and an analysis software NIBPCHART (Panlab, #76-0466,) early in the morning, and the median of three consecutive measurements was used as the recorded value. Using the blood pressure measured in the morning before mating as the baseline, measurements were taken daily from E0.5 to E18.5 after mating and plug confirmation. Variations relative to the prepregnancy blood pressure were subsequently analyzed.

### Neonatal follow-up

Neonatal weights were measured weekly at postnatal week 1, 2, 3 and 4. Neuromotor behavior was examined on the morning of postnatal day 5 using the surface righting test. In this test, each infant was placed on its back on a flat table, and the time taken by the infant to get to the supine position was measured (maximum 60 s) ^67^.

### Statistical analyses

Statistical analyses were conducted using GraphPad Prism Version 10 (GraphPad Software). Outliers were identified and removed. Normality was evaluated using the Shapiro-Wilk test. To determine the significant differences, the Mann–Whitney U test was used for nonparametric data, unpaired two-tailed Student’s *t*-test was used for parametric data, and Fisher’s exact test was used for binary data. One-way analysis of variance followed by Dunnett’s post hoc test was used for comparison of data between multiple groups. The Wilcoxon signed-rank test was used as the nonparametric test for paired samples.

## Supplemental Figures

**Figure S1.**
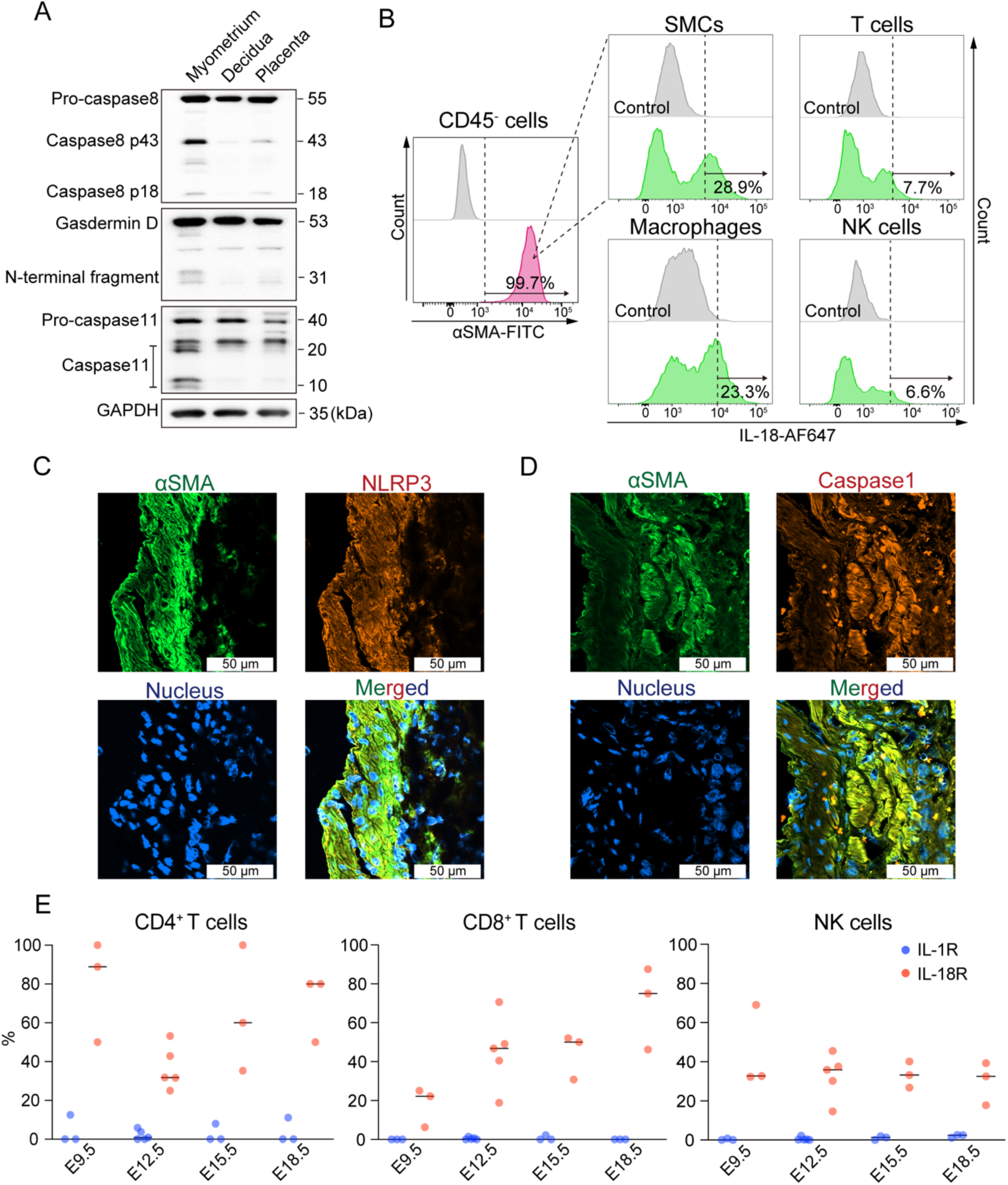
IL-18 is produced in uterine myometrium, related to Figure 1. (A) Immunoblotting using the indicated antibodies. Uterine myometrium, decidua and placenta were extracted from pregnant mice, and the cells were collected from whole tissue lysates. The image is representative of independent two samples. (B) Representative histogram of flow cytometric analysis showing APC-labeled IL-18 in SMCs, T cells, macrophages, NK cells of uterine myometrium. Each image is representative of independent two samples. (C, D) Confocal fluorescent immunohistochemistry images using indicated antibodies. Pregnant uteri were extracted at E9.5. Each image is representative of independent two samples. (E) Cell surface positivity rates of IL-1R and IL-18R on each immune cell populations are shown by each gestational stage (each data point represents 1 dam, n = 3–5 per each group, each horizontal line represents mean value of samples).

**Figure S2.**
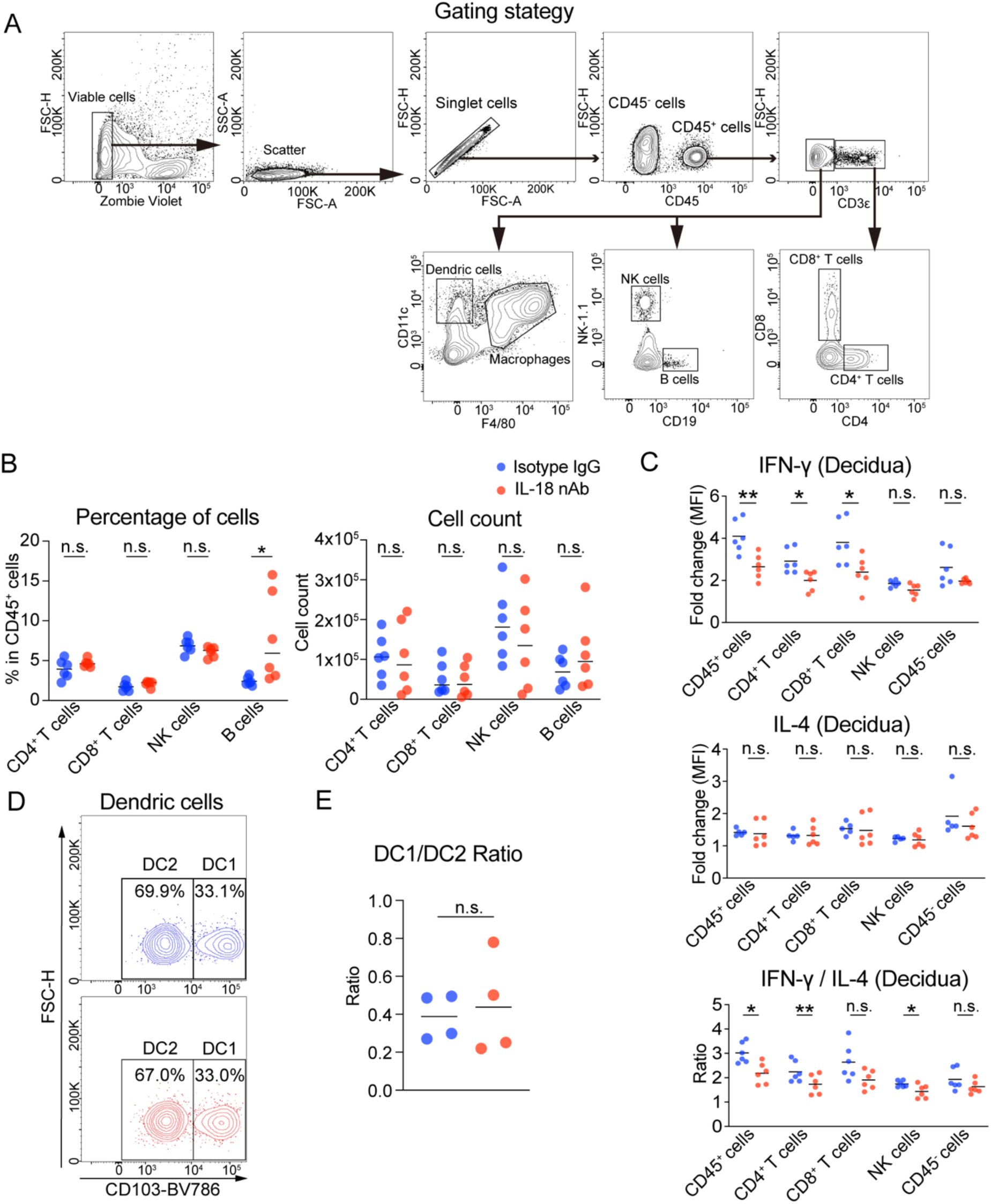
IL-18 promotes type 1 immune responses in uterus, related to Figure 2. (A) Multicolor flow cytometric gating strategy is shown. (B) The percentage of each cell in CD45^+^ cells in both groups is shown on the left, and the cell count of each cell in both groups is shown on the right (Mann-Whitney U test, each data point represents 1 dam, n = 6 per each group, each horizontal line represents mean value of samples). (C) Mean fluorescence intensity (MFI) of PE-Cy7-labeled intracellular cytokines in each decidual immune cell populations is shown for relative change to control by group (Mann-Whitney U test, each data point represents 1 dam, n = 6 per each group, each horizontal line represents mean value of samples). (D) Representative contour plots of flow cytometric analysis showing distribution of DC1 (F4/80^-^ CD11c^+^CD103^+^) and DC2 (F4/80^-^CD11c^+^CD103^-^) dendric cells identified by Brilliant Violet (BV) 786- labeled CD103. (E) The ratios of DC1/DC2 cell population in dendric cells are shown by group (Student’s *t*-test, each data point represents 1 dam, n = 4 per each group, each horizontal line represents mean value of samples). * *p* < 0.05, ** *p* < 0.01

**Figure S3.**
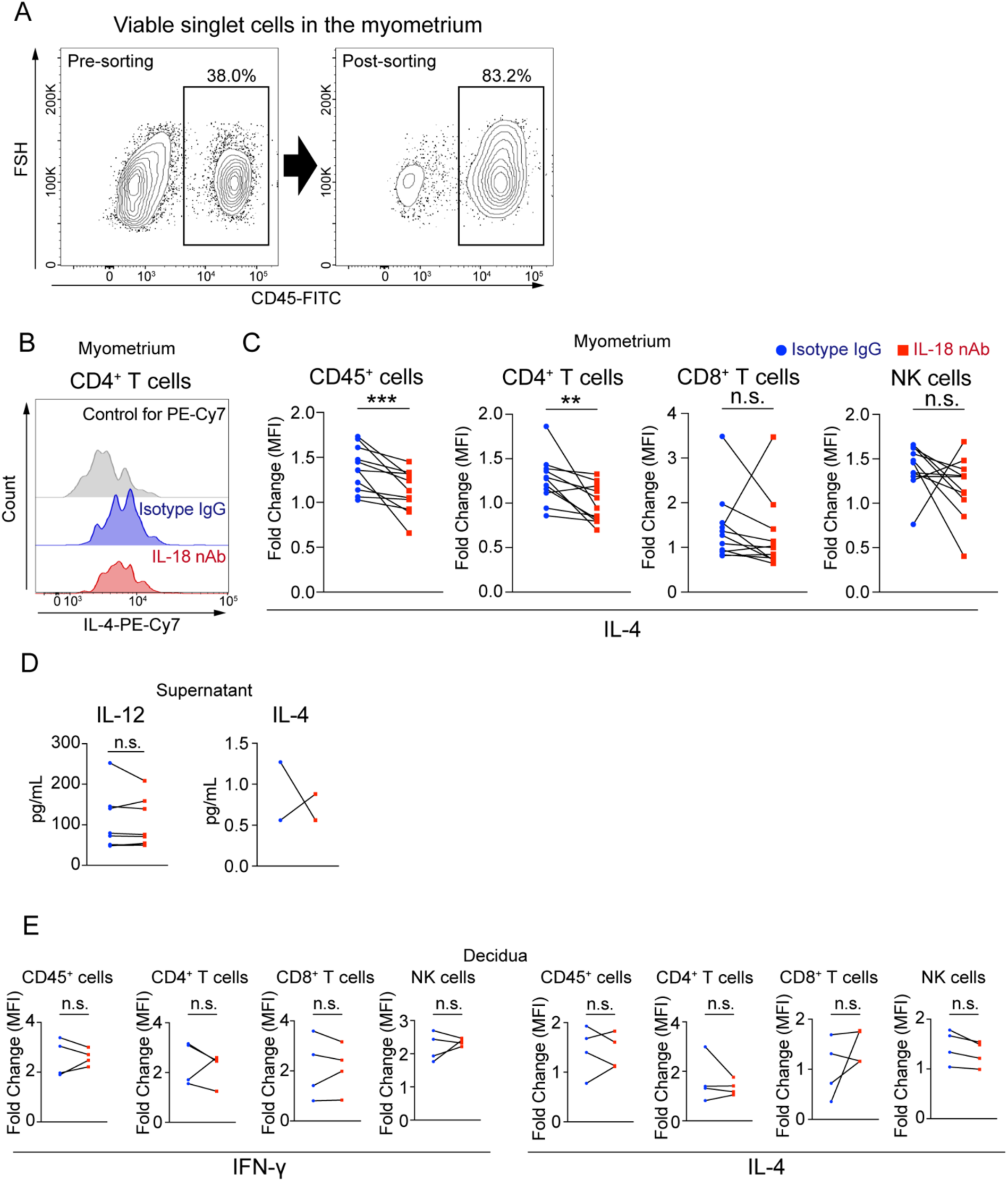
IL-18 also promotes IL-4 production in uterus *in vitro*, related to Figure 3. (A) Representative contour plots of flow cytometric analysis showing purity of cell sorting with MACS identified by BV510-labeled CD45. (B) Representative histogram of flow cytometric analysis showing PE-Cy7-labeled IL-4 in CD4^+^ T cells by group. (C) MFI of PE-Cy7-labeled intracellular IL-4 in each immune cell populations is shown for relative change to control by group (Wilcoxon matched-pairs signed rank test, each pair of data points represents 1 dam, n = 11 per each group). (D) The concentration of IL-12 and IL-4 in culture supernatant quantified with ELISA is shown (Wilcoxon matched-pairs signed rank test, each pair of data points represents 1 dam, n = 9 per each group, seven samples including values below the detection sensitivity leads to the removal of the corresponding plots in the IL-4 measurement). (E) MFI of PE-Cy7-labeled intracellular IFN-γ and IL-4 in each decidual immune cell populations is shown for relative change to control by group (Wilcoxon matched-pairs signed rank test, each pair of data points represents 1 dam, n = 4 per each group). ** *p* < 0.01, *** *p* < 0.001

**Figure S4.**
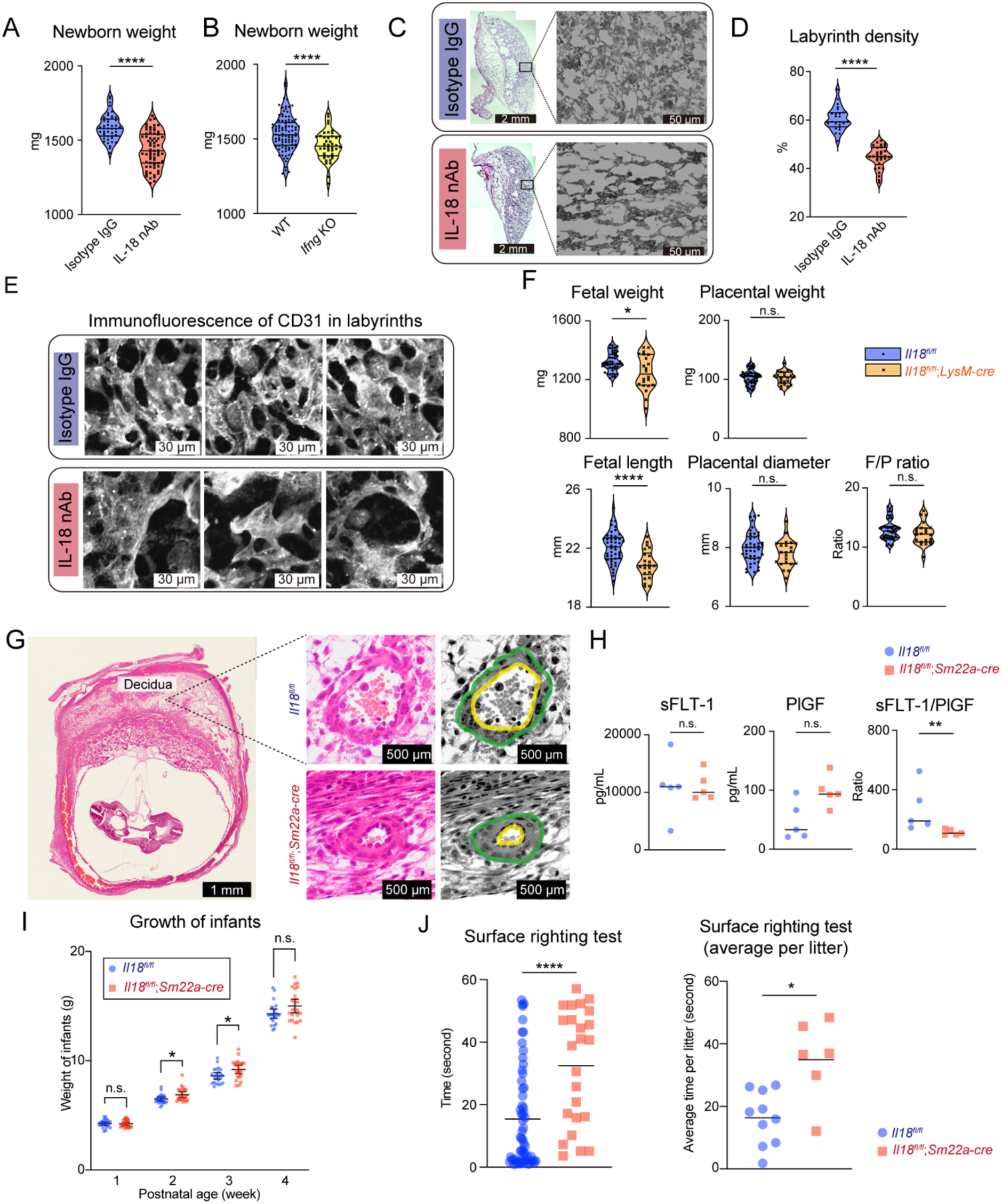
IL-18-IFN-γ axis plays critical role in fetoplacental growth, related to Figure 4. (A, B) Violin plots representing newborn weight after spontaneous birth by group (Student’s *t*-test, each data point represents 1 newborn, 3–5 dams per each group, each horizontal line represents median and first and third quartile of samples). (C) Representative frozen section of E9.5 placenta with H&E staining on left. Representative gray-scaled high-power field pictures in placental labyrinth area with H&E staining on right. (D) Violin plots representing density of labyrinths quantified from binarized HE stained images. (Student’s *t*-test, each data point represents a mean value of three regions of interest (ROI), three ROI was randomly selected from labyrinth in each section, 24–27 sections from three pregnancies by each group, each horizontal line represents median and first and third quartile of samples). (E) Confocal fluorescent immunohistochemistry images of frozen sections of labyrinth using CD31 antibody by group. Pregnant uteri were extracted at E9.5. (F) Violin plots representing fetal weight, placental weight, fetal length and placental diameter at E18.5 are shown by group. (Mann-Whitney U test, each data point represents 1 newborn or placenta, 3–4 dams per each group, each horizontal line represents median and first and third quartile of samples). (G) Representative section of E9.5 implantation site of pregnant mice with H&E staining on left. Representative high-power field pictures show stereological assessment of wall (green line) and lumen (yellow line) of uterine spiral arteries on right. (H) The concentration of sFLT-1 and PlGF in the serum collected from pregnant mice at E13.5 was quantified with ELISA and shown by group. The ratio of sFLT-1 to PlGF is also shown by each group (Mann-Whitney U test, each data point represents 1 dam, n = 5 per each group, each horizontal line represents mean value of samples). (I) Weights of infants born by spontaneous delivery were measured at 1, 2, 3, and 4 weeks postpartum and are shown for each group (Mann-Whitney U test, each data point represents 1 infant, n = 22–24 from 4 dams per each group, each horizontal line represents mean value of samples and ±standard error of the mean). (J) Neuro-motor behavior was examined with surface righting test. The time it took for each neonate to return from the supine to the prone position (lest) and the mean value of that time for each litter (right) are shown for each group (Mann-Whitney U test, each data point represents 1 infant in the left and the mean value of each litter in the right, 2–8 litters from 6–8 dams per each group, each horizontal line represents mean value of plots) * *p* < 0.05, ** *p* < 0.01, **** *p* < 0.0001

**Figure S5.**
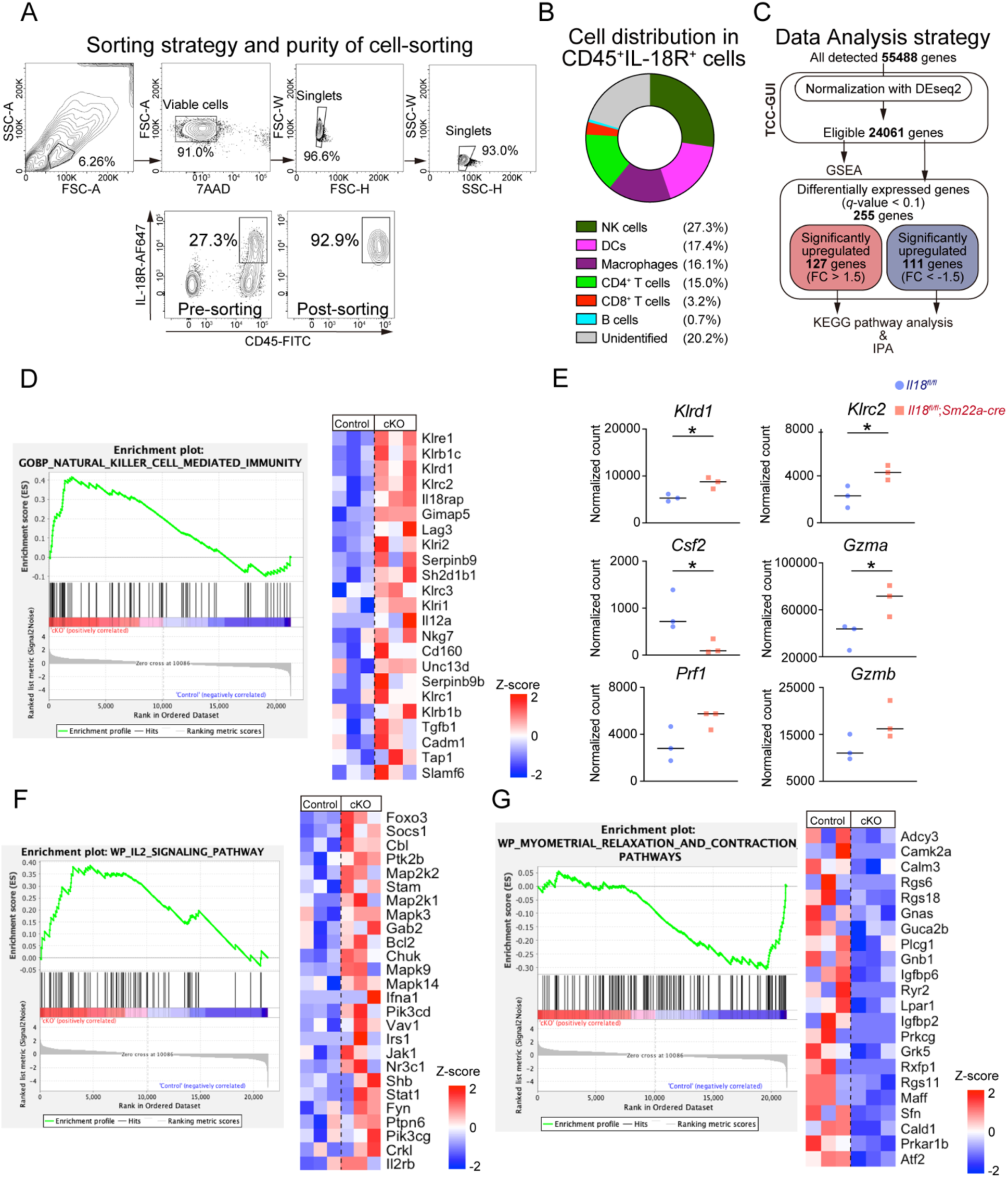
RNA-seq reveals impact of IL-18-signaling on the immune milieu, related to Figure 5. (A) Multicolor flow cytometric sorting strategy and its purity is shown. (B) The breakdown of cells enriched by cell sorting is shown in the doughnut chart. (C) The flowchart shows the data analysis strategy for RNA-seq. Volcano plots depicting the results of the RNA-seq study with highlighted plots representing significantly upregulated or downregulated genes. (D, F, G) The enrichment plot for each pathway analyzed by GSEA is shown on the left, and the heatmap from the Z-score of the significantly enriched genes among the pathway’s gene set is shown on the right. (Related to Table S4) (E) For each gene involved in NK cell function, normalized counts identified by RNA-seq were plotted (Student’s *t*-test, each data point represents 1 sample, n = 3 per each group, each horizontal line represents mean value of samples). * *p* < 0.05

**Figure S6.**
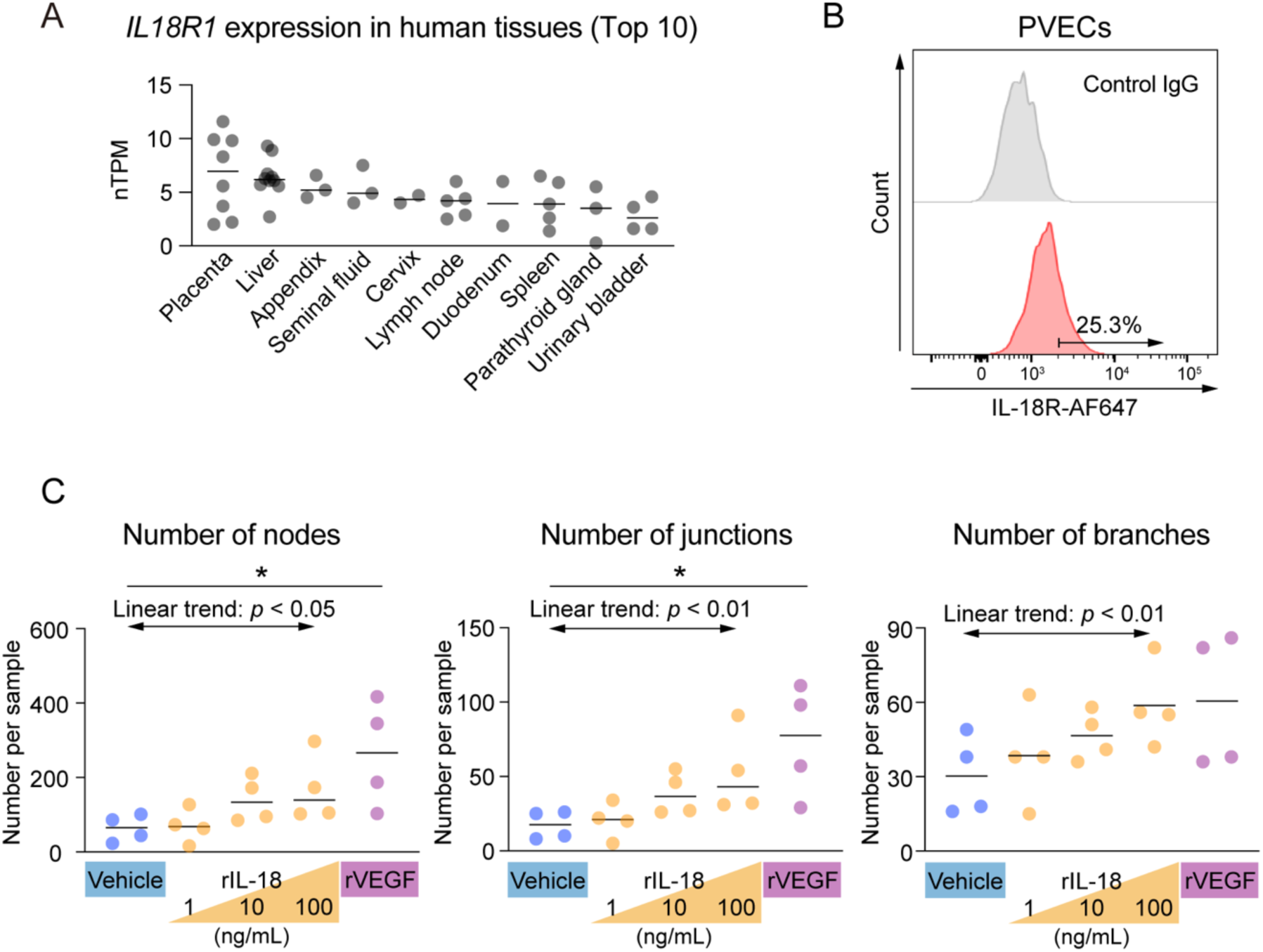
IL-18 promotes placental angiogenesis *in vitro*, related to Figure 6. (A) Gene expression of *IL18R1*, which encodes the IL-18R, was cited from the HPA RNA-seq database (data available from https://www.proteinatlas.org/ENSG00000115604-IL18R1/summary/rna) and sorted by expression level in each systemic tissue and shown as top 10. (n = 2–10 per each group) (B) Representative histogram of flow cytometric analysis showing AF647-labeled IL-18R on PVECs. The image is representative of two samples. (C) Images obtained by tube formation assay were quantified using the Angiogenesis Analyzer for ImageJ and plotted for each indicated parameter by each treatment (One-way analysis of variance, linear trends from vehicle to 100 ng/mL rIL-18 treatment were also analyzed, each data point represents 1 sample, n = 4 per each group, each horizontal line represents mean value of samples). * *p* < 0.05

## Notes

### Competing Interest Statement

The authors have declared no competing interest.

